# MoHuTS: Developability-Aware Antibody Humanization Using Multi-Objective Tree Search

**DOI:** 10.64898/2026.11.24.768826

**Authors:** Kiichi Yoshimura, Satoru Fujii, Shoichi Ishida, Kei Terayama

## Abstract

Antibody humanization aims to reduce the immunogenicity risk associated with non-human antibodies while preserving antigen binding and other properties important for therapeutic development. Identifying suitable framework substitutions often requires repeated rounds of variant construction and experimental testing, making humanization costly. Recently, machine-learning methods, including language models trained on antibody sequences, have been proposed to identify human-like framework substitutions. Although prior humanization methods have considered antigen binding and biophysical properties, it remains challenging to balance humanness across both chains with developability-related properties such as thermal stability and aggregation propensity while retaining favorable parental properties. Here, we present Multi-objective Humanization Tree Search (MoHuTS), which combines an antibody language model with reward-guided tree search to explore combinations of framework substitutions efficiently. Starting from the parental heavy- and light-chain sequences, the language model proposes successive framework substitutions. A combined score for humanness, melting temperature, a structural proxy for binding retention, and aggregation propensity guides the search. Across five computational case studies, MoHuTS generated candidates that showed favorable humanness scores and developability-related properties with relatively few framework substitutions from the parental sequences. These results suggest that MoHuTS can help prioritize humanization candidates for experimental evaluation.

## 1. Introduction

A key challenge in antibody humanization is to reduce the immunogenicity risk associated with non-human antibodies while preserving antigen binding and developability related properties required for therapeutic development (Choi et al., 2016; G. L. Gordon et al., 2024). Although fully human antibodies can be obtained from transgenic mice or human antibody phage-display libraries (Lonberg et al., 1994; Marks et al., 1991), humanization remains a widely used route for developing therapeutic antibodies from non-human precursors (G. L. Gordon et al., 2024; Lu et al., 2025). A conventional approach grafts the parental complementarity-determining regions (CDRs) onto human variable-region frameworks (Riechmann et al., 1988). However, substitutions at certain framework positions can affect antigen-binding affinity, thermal stability, aggregation propensity and other biophysical properties relevant to developability even if the CDR sequences are unchanged (Foote and Winter, 1992; Gupta et al., 2024). Even when antigen binding is retained, thermal stability and aggregation propensity are important considerations when selecting candidates for further development (Jain et al., 2017). Some framework residues may therefore be reverted to their parental identities through back mutation to recover properties (Kettleborough et al., 1991; G. L. Gordon et al., 2024). Consequently, identifying a humanized sequence that retains antigen binding and favorable developability related properties can require repeated rounds of variant construction and experimental characterization (Tennenhouse et al., 2024).

Machine-learning methods have been developed to reduce this experimental burden. For example, Hu-mAb (Marks et al., 2021) iteratively selects framework substitutions that maximize a variable (V)-gene-specific humanness score until the score reaches a criteria, aiming to limit substitutions. Sapiens (Prihoda et al., 2022), a language model trained on human antibody sequences, proposes framework substitutions using context-dependent amino acid probabilities. Humatch (Chinery et al., 2024) combines V-gene-specific convolutional neural networks, a classifier for heavy-chain variable domain (VH) and light-chain variable domain (VL) pairing, and germline sequence information to guide joint humanization of heavy and light chains. However, these methods primarily optimize sequence humanness without explicitly considering antigen-binding affinity or biophysical properties.

To account for important properties other than the degree of humanization, several methods have been reported that partially incorporate developability related properties into the humanization process. For example, computationally driven antibody humanization (CoDAH) selects combinations of framework substitutions by balancing humanness with structure-based energy scores (Choi et al., 2015, 2016). Computational human antibody design (CUMAb) ranks human-framework grafts using Rosetta energy and filters candidates for CDR backbone preservation (Tennenhouse et al., 2024). Its search focuses on alternative human frameworks rather than exploring individual framework substitutions around the parental sequence. However, neither method explicitly uses predicted melting temperature (*T_m_*) or aggregation propensity as a separate optimization objective. Gordon et al. demonstrated property-guided humanization using affinity and thermostability predictors for a variable domain of a heavy-chain-only antibody (VHH), but did not evaluate paired VH/VL sequences (C. Gordon et al., 2024). Hu-MCTs optimizes humanness and CDR structural preservation in paired antibodies, but its reported candidates contain extensive changes from the parental sequences, and its objectives do not include thermostability or aggregation propensity (Chen et al., 2026). Despite these advances in antibody humanization, there is still room to develop methods that improve the humanness of VH and VL sequences while preserving antigen binding and accounting for key properties relevant to developability, including predicted melting temperature (*T_m_*) and aggregation propensity. In addition, jointly optimizing humanness and these properties with as few framework substitutions as possible could help preserve the favorable biophysical properties of the parental antibody.

To this end, we propose Multi-objective Humanization Tree Search (MoHuTS), a developability-aware method that uses a modified upper confidence bounds applied to trees (UCT) rule (Kocsis and Szepesvári, 2006) to efficiently explore combinations of framework substitutions starting from the parental VH/VL pair. MoHuTS generates candidate variants by applying human-like framework substitutions proposed by the antibody language model Sapiens (Prihoda et al., 2022), with the search implemented using ChemTSv3 (Fujii et al., 2026). The search is guided by a reward that integrates four scores for sequence humanness assessed by Observed Antibody Space identity search (OASis) (Prihoda et al., 2022), predicted thermal stability, preservation of modeled CDR conformations as a proxy for binding retention (Zhang et al., 2013; Apgar et al., 2016), and aggregation propensity estimated from the AGGRESCAN scale (Conchillo-Solé et al., 2007). Across five computational case studies, MoHuTS generated candidates with favorable humanness scores and predicted biophysical properties with relatively few changes to the parental framework sequences. In additional analyses, high-reward candidates had higher mean human germline framework identity and AbNatiV scores (Ramon et al., 2024) than their parental sequences. We also evaluated modeled antibody– antigen interfaces using InterfaceAnalyzer (Stranges and Kuhlman, 2013) in Rosetta (Leaver-Fay et al., 2011) and found that several interface metrics were close to or more favorable than the corresponding parental values. These results suggest that MoHuTS could support the prioritization of humanization candidates for experimental evaluation.

## 2. Methods

### 2.1 Study design and overall workflow

MoHuTS uses UCT (Kocsis and Szepesvári, 2006) to search for antibody candidates that balance humanness and predicted biophysical properties (Fig. 1(A)). The search was represented as a tree rooted at the parental VH/VL sequence pair, with each node representing a candidate pair. Each edge represented a single framework substitution in either VH or VL, while Kabat-defined CDRs (Kabat et al., 1991) remained unchanged. Substitutions were proposed using Sapiens, a language model trained on human antibody sequences (Prihoda et al., 2022). As shown in Fig. 1(B), the search comprises four steps: selection, expansion, evaluation, and backpropagation. Selection balanced further search along paths that had produced high-scoring candidates with exploration of less-visited paths. During expansion, the selected VH and VL sequences were given to Sapiens to propose framework substitutions, and the resulting pairs were added as child nodes. Each candidate was evaluated for humanness, predicted thermal stability, CDR structural preservation, and predicted aggregation propensity. These four scores were combined into a single integrated reward, which was backpropagated along the selected path to the root, updating node statistics for subsequent selection. By repeating these steps, MoHuTS explored candidates that balanced humanness and predicted biophysical properties.

**Figure 1.**
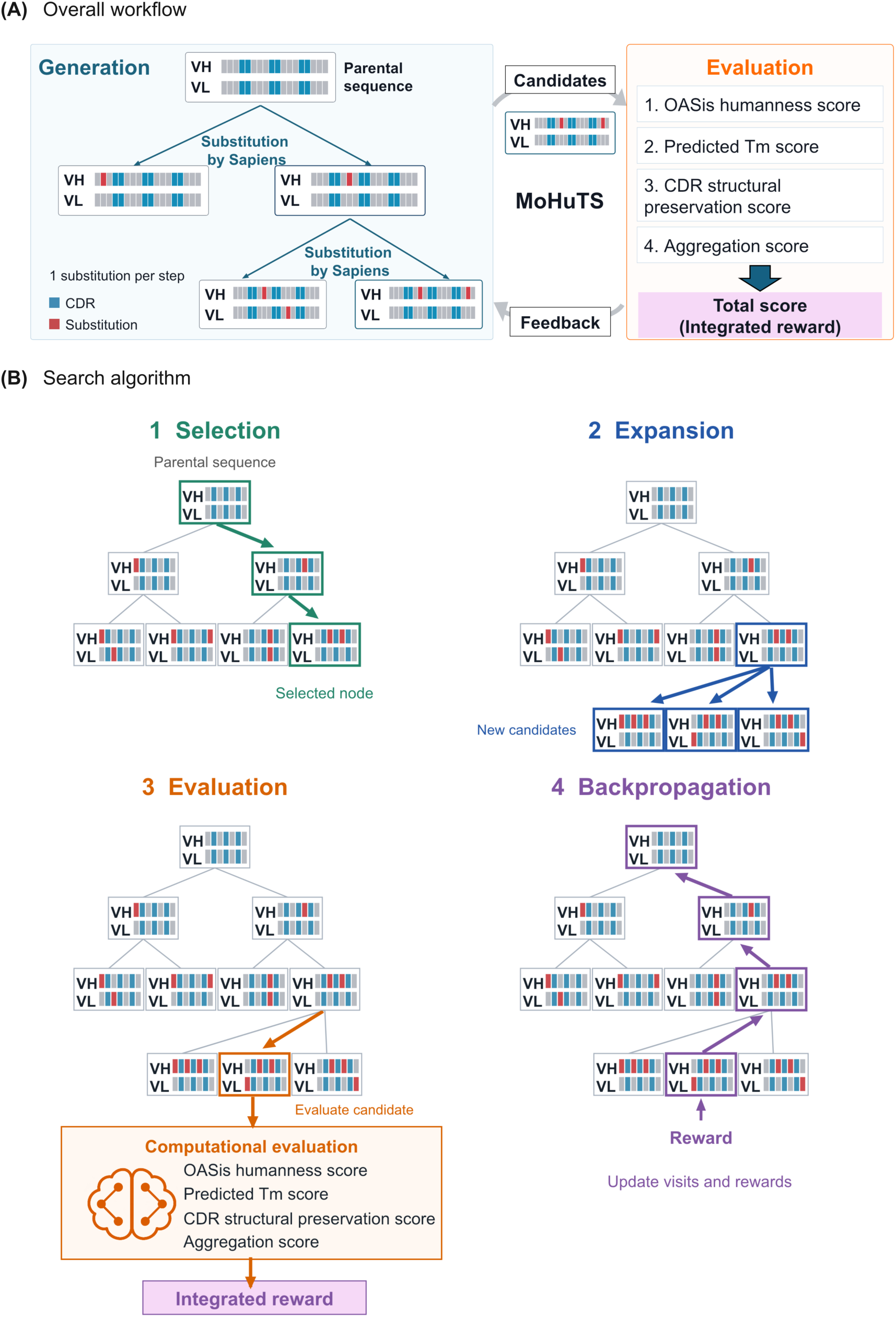
Overview of MoHuTS. Panel A shows the workflow, starting from the paired parental VH and VL sequences at the top. Each node represents a candidate VH/VL pair; each edge represents a single VH or VL framework substitution. UCT guides selection along existing branches. Sapiens is applied to the current VH and VL sequences at each node selected for expansion. Red blocks indicate substitutions from the parent, and cyan blocks indicate unchanged Kabat-defined CDRs. OASis humanness, predicted *T*_m_, CDR structural preservation, and aggregation scores form an integrated reward, which is backpropagated along the selected path (purple arrows) to guide selection. Candidates are ranked by reward after the search. Panel B illustrates the repeated steps of UCT-guided search: selection, expansion, evaluation, and backpropagation.

### 2.2 CDR definition and mutation constraints

CDRs were excluded from mutation to reduce the risk of impairing antigen binding. We assigned Kabat numbering to each VH and VL sequence using AbNumber, which uses the antigen receptor numbering and receptor classification tool (ANARCI; Dunbar and Deane, 2016). CDRs comprised VH positions 31–35, 50–65, and 95–102, and VL positions 24–34, 50–56, and 89–97 (Kabat et al., 1991). Residues with insertion codes were included if the numeric part of their Kabat number fell within these CDR ranges; thus, VH positions 35A and 35B were excluded from mutation.

### 2.3 Sapiens-guided mutation proposal

Framework substitutions were proposed using Sapiens, which predicts a probability distribution over the 20 standard amino acids at each position in the current VH or VL sequence based on human antibody sequence patterns (Prihoda et al., 2022). A lower probability for the current amino acid indicates that it is less consistent with human antibody sequence patterns at that position. During tree search, we applied Sapiens separately to VH and VL at each node selected for expansion. For each chain, we selected up to 16 framework positions with the lowest probabilities for the current amino acids. At each selected position, we replaced the current residue with the most probable alternative to obtain a single-substitution candidate. This produced up to 32 candidate VH/VL sequences as child nodes per expansion.

### 2.4 UCT-guided tree search

The tree search used a modified UCT rule to balance high-scoring and less-visited branches, implemented using ChemTSv3, a framework for molecular generation (Fujii et al., 2026). Selection prioritized unvisited children, choosing one uniformly at random. Once all children had been visited, the child with the highest modified UCT score was selected. For child *i*, the score is:

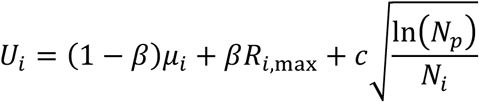

where *μ_i_* and *R_i_*_,max_ are the mean and maximum integrated rewards backpropagated through child *i*, respectively, and *N_p_* and *N_i_* are the parent and child visit counts. The parameter *β* balances mean and maximum rewards, while *c* scales exploration of less-visited children. All five antibody cases used *c* = 0.01 and *β* = 0.5. At each expansion, proposed VH/VL sequences were added as children of the selected node. One child was evaluated; the others remained available for later selection. Its integrated reward was backpropagated along the selected path to update visit counts, mean rewards, and maximum rewards.

### 2.5 Calculation and integration of four-component scores

Sequence humanness was evaluated using the OASis_9mers_v1 database at a prevalence threshold of 10% of human subjects (Prihoda et al., 2022). The paired-chain OASis score was the number of qualifying 9-mer windows across both chains divided by the total number of windows. Windows were counted separately even when the same 9-mer occurred at multiple positions. This 0–1 score served directly as the humanness component of the reward.

We developed a regression model to predict melting temperature (*T*_m_). Each VH/VL sequence was represented as a 480-dimensional vector using AbLang-2 (Olsen et al., 2024). These vectors were used to predict melting temperature (*T*_m_) with a LightGBM (Ke et al., 2017) model trained on 165 antibody sequences with experimentally measured melting temperatures (Jain et al., 2017; Shehata et al., 2019). Five-fold cross-validation yielded a Pearson correlation coefficient of *r* = 0.660 and a root-mean-square error (RMSE) of 5.024 °C (training and evaluation details in Supplementary Methods S1). Predicted melting temperature, *T*_m_ (°C), was converted into a normalized score using a sigmoid function:

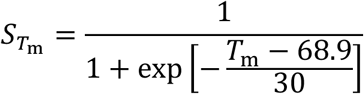

The midpoint of 68.9 °C is the training dataset’s mean experimental *T*_m_ and maps to a score of 0.5. The scale parameter of 30 °C controls how gradually scores change with temperature; higher predicted temperatures yield higher scores.

To assess potential effects of framework substitutions on antigen binding, we measured CDR backbone deviations from the parental antibody, consistent with previous structure-guided humanization studies (Zhang et al., 2013; Apgar et al., 2016). Parental and candidate VH/VL structures were predicted using ABodyBuilder2 (Abanades et al., 2023). Each candidate was aligned with its parent by a single rigid-body fit to corresponding framework backbone atoms (N, CA, C, and O) from both chains. We then calculated backbone root-mean-square deviations (RMSDs) between corresponding CDR loops without separately realigning them. RMSDs of all loops among the six CDRs were averaged with equal weight to obtain the mean CDR RMSD, *d* (Å), and normalized using a decreasing sigmoid:

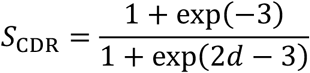

The numerator ensures a score of 1 at *d* = 0; scores decrease as mean CDR RMSD increases. The coefficient of 2 controls steepness, and the offset of 3 sets the inflection point at *d* = 1.5 Å, where the score is approximately 0.525.

The aggregation score used the AGGRESCAN scale of amino acid aggregation-propensity values (A3V; Conchillo-Solé et al., 2007), favoring lower VH aggregation propensity while penalizing deterioration in the length-weighted VH/VL value. Residue-level A3V values were averaged over a seven-residue sliding window to obtain each chain’s average aggregation-propensity profile (A4V). This profile was averaged across the chain and multiplied by 100 to obtain its raw aggregation value, *N_c_*; lower values indicate lower sequence-based aggregation propensity. Chain values were combined using a sequence-length-weighted mean:

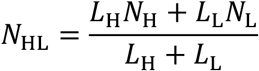

Here, *N*_H_ and *N*_L_ are the raw VH and VL values, and *L*_H_ and *L*_L_ are their respective residue counts. Candidate-minus-parental differences in VH and paired-chain values, Δ*N*_H_ and Δ*N*_HL_, were converted to an aggregation score:

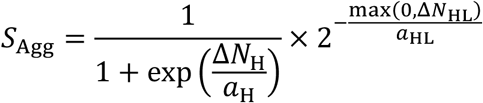

The first factor increases as the VH raw value decreases relative to the parent. The second equals 1 when the paired-chain value is unchanged or decreases and falls below 1 when it increases. The score was clipped to [10⁻⁶, 1 − 10⁻⁶] before reward integration; the parent scores 0.5. Scale constants *a*_H_ and *a*_HL_ were derived from 100 VH/VL sequences: parental, experimentally validated humanized, Sapiens-generated, and Hu-mAb-generated sequences for each of 25 antibodies. We set *a*_H_ = 5.211315 and *a*_HL_ = 3.973924 to the interquartile ranges (IQRs; 75th minus 25th percentile) of raw VH and paired-chain values, respectively. These constants control the effect of parent-relative changes on the score.

The four-component scores were combined by a weighted geometric mean. Each ranged from 0 to 1, with higher values indicating more favorable properties. For a VH/VL sequence *x*, the integrated reward was:

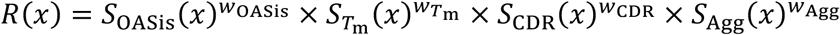

Here, *S*_OASis_(*x*), *S_T_*_m_(*x*), *S*_CDR_(*x*), and *S*_Agg_(*x*) are the component scores for humanness, predicted thermal stability, CDR structural preservation, and aggregation propensity, respectively. The weighted geometric mean reduces the reward when a component score is low, encouraging balance across objectives. All five antibody cases used *w*_OASis_ = 0.4, *w_T_*_m_ = 0.2, *w*_CDR_=0.1, and *w*_Agg_ = 0.3. Humanness received the largest weight to prioritize humanization. The aggregation score received the largest weight among the biophysical components because protein aggregation is a developability concern and may increase immunogenicity risk (Joubert et al., 2012; Rombach-Riegraf et al., 2014). The *T*_m_ score received an intermediate weight to favor thermal stability while limiting the influence of a model with moderate predictive accuracy. CDR structural preservation received the smallest weight because CDR sequences were already fixed, and this score provided an additional check for framework-induced CDR structural changes.

### 2.6 Target antigens and search settings

We evaluated MoHuTS in five computational humanization cases corresponding to certolizumab, pertuzumab, bevacizumab, omalizumab, and eculizumab. Their respective targets were tumor necrosis factor α (TNF-α), human epidermal growth factor receptor 2 (HER2), vascular endothelial growth factor A (VEGF-A), immunoglobulin E (IgE), and complement component 5 (C5). Starting from the corresponding parental VH/VL sequences (Data S1), we performed five independent searches with different random seeds per parent. Each stopped after exploring 3,000 distinct VH/VL sequences.

### 2.7 Antibody–antigen interface analysis

Rosetta (Leaver-Fay et al., 2011) was used to evaluate the antibody–antigen interfaces of the five highest-reward candidates and reference sequences after candidate generation; its scores did not contribute to the reward. For each sequence, Rosetta Relax generated one relaxed structure using the ref2015 score function (Alford et al., 2017) and starting-structure coordinate restraints, while allowing backbone movement. InterfaceAnalyzer (Stranges and Kuhlman, 2013) evaluated the relaxed structure using the same seed and score function, with side-chain repacking before and after antibody–antigen separation. Each variant’s interface score *dG* was summarized by the median of three completed runs. Subtracting the parental median yielded the parent-relative difference:

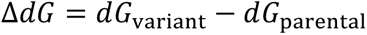

Negative differences in Rosetta energy units (REU) indicate lower modeled interface energies than the parent. Table S2 provides template structures, chain assignments, and interface scores.

## 3. Results and discussion

### 3.1 Substitution search using MoHuTS in five antibody humanization cases

Starting from parental VH/VL sequences, MoHuTS searched for candidates balancing humanness and predicted biophysical properties in five humanization cases. Mean integrated reward (red line) increased in each search (Fig. 2A–E), reaching approximately 0.67–0.73 and exceeding parental, experimentally validated humanized, Sapiens, and Hu-mAb reference rewards. Since CDR structural preservation was assessed relative to the parent and parental aggregation scores were generally higher, the parental sequences (black lines) and experimentally humanized references (orange dashed lines) had similar integrated rewards in some cases.

**Figure 2.**
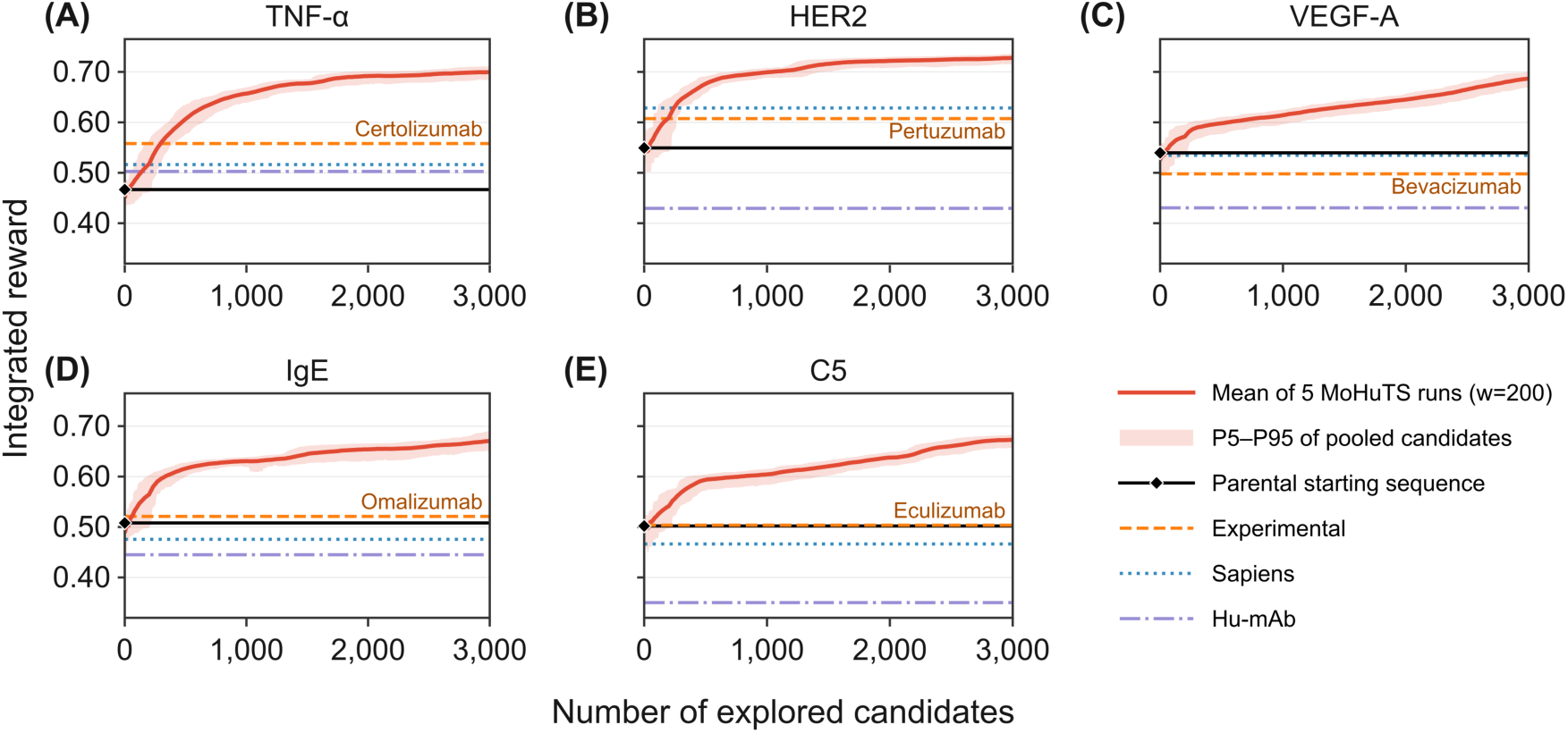
Integrated-reward trajectories across five antibody humanization cases. Panels A–E show antibodies targeting TNF-α, HER2, VEGF-A, IgE, and C5, respectively (n = 5 MoHuTS runs per case). Each run explored 3,000 unique VH/VL sequences. Solid red lines show trailing 200-candidate means averaged across the five runs. Shading spans the 5th–95th percentiles of rewards pooled across runs within the same windows, showing variation among candidates. Black diamonds mark parental rewards at x = 0. Horizontal lines indicate parental, experimentally validated humanized (orange dashed lines labeled by antibody name), Sapiens, and Hu-mAb references. The x-axis shows explored candidates per run.

Figure 3A–E shows component-score trajectories for TNF-α; Fig. 3F, G shows VH and VL alignments of the top-five candidates and references, discussed in detail later. Integrated reward increased overall alongside humanness, predicted *T_m_*, and aggregation scores. Mean CDR structural preservation declined but remained high; the five highest-reward candidates scored 0.949–0.965 versus 0.991 for the parent (Table S3). The other four cases showed similar overall increases in mean reward, humanness, and aggregation scores, with declining but high CDR structural preservation scores (Fig. S1). Experimental references had lower CDR structural preservation and aggregation scores than parents in all five cases (Fig. 3D, E and Fig. S1), contributing to rewards close to or below parental values in some cases (Fig. 2A–E).

**Figure 3.**
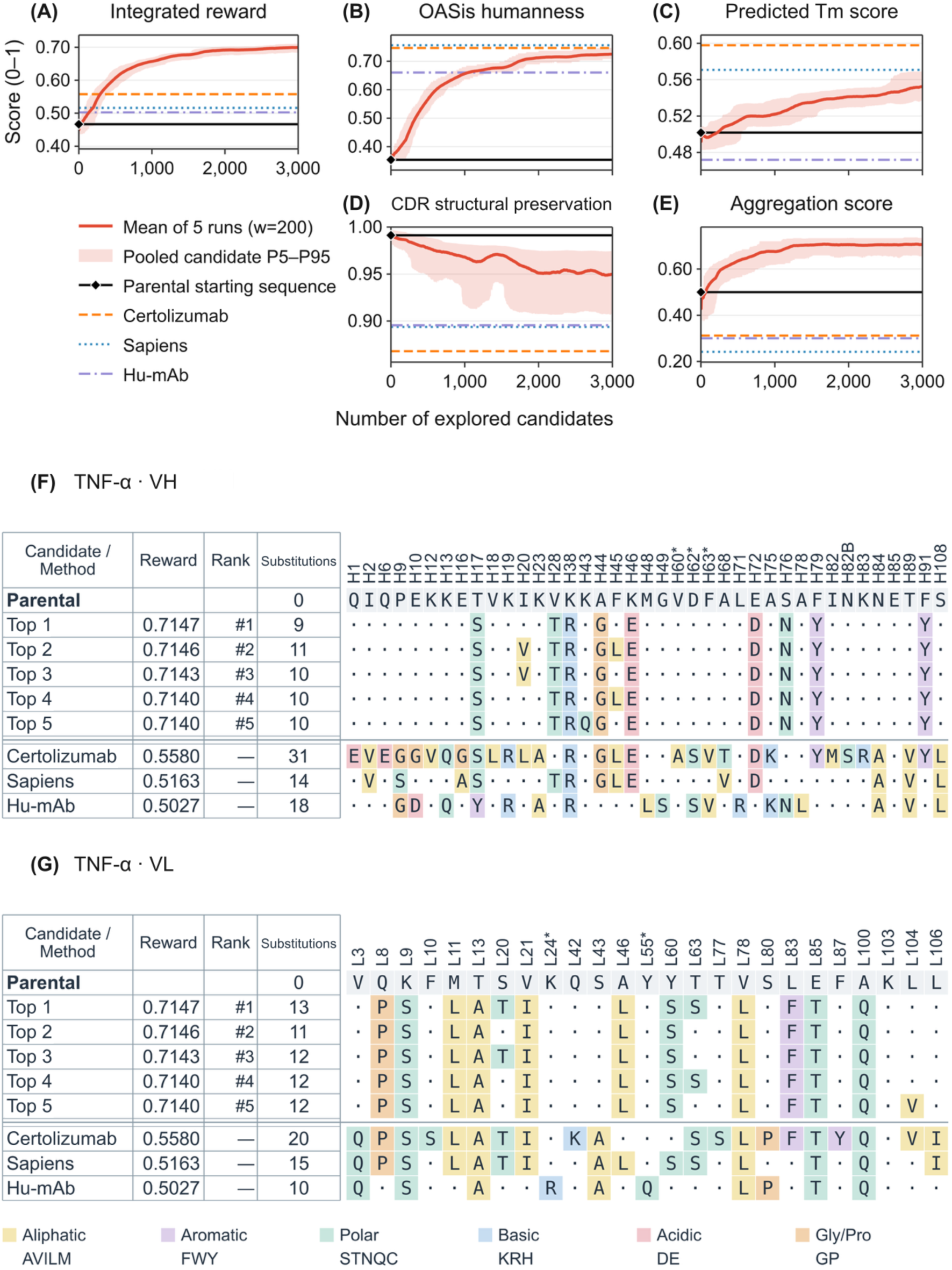
Search trajectories and sequence changes during anti-TNF-α antibody humanization. Panels A–E show integrated reward, OASis humanness, predicted Tm score, CDR structural preservation, and aggregation scores, respectively. Solid red lines show trailing 200-candidate means across five independent runs. Shading spans the 5th– 95th percentiles of scores pooled across runs within the same windows, showing variation among candidates. Black diamonds mark parental scores at x = 0. Horizontal lines indicate parental, certolizumab (orange dashed line), Sapiens, and Hu-mAb references. The x-axis shows unique candidates explored per run. Panels F and G show VH and VL alignments, respectively, of the top-five distinct VH/VL sequences (Top 1–Top 5, ranked by reward) and references, with all parent-relative substitutions. Reward, rank, and top-five summaries refer to paired VH/VL sequences; substitution counts are parent-relative for each chain, and comparators are unranked. Dots denote parental residues, asterisks mark Kabat CDR positions, and colors indicate amino acid classes.

We visualized sequence changes during searches using principal component analysis (PCA) (Fig. 4A–E). Each VH/VL sequence was represented numerically using AbLang-2 (Olsen et al., 2024), and PCA summarized the main variation in two dimensions. Across all five cases, later candidates generally appeared closer to experimentally validated humanized or Sapiens references than earlier candidates, although the extent varied.

**Figure 4.**
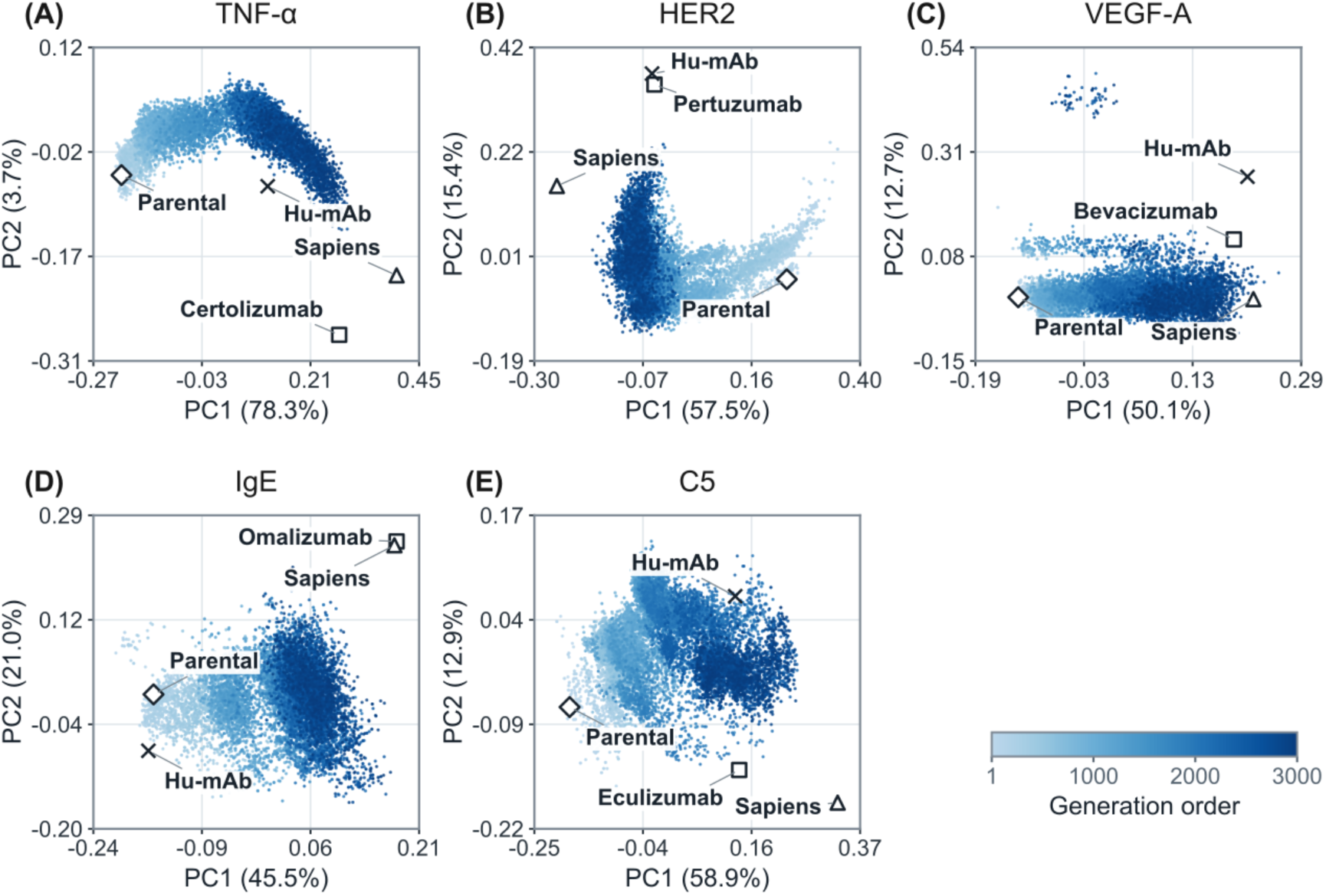
PCA of generated and reference sequence features. Panels A–E show antibodies targeting TNF-α, HER2, VEGF-A, IgE, and C5, respectively, each with 15,000 candidate records from five runs (3,000 per run). Points are colored by generation order within runs, from light blue (earlier) to dark blue (later). PCA summarizes AbLang-2 sequence features in two dimensions. For each antibody, axes were determined using candidates from all five runs. Parental, experimentally validated humanized (open squares labeled by antibody name), Sapiens, and Hu-mAb references were plotted on the same axes. Axis labels show the variance percentage explained by each component.

### 3.2 Detailed characterization of the highest-reward MoHuTS candidates

The five highest-reward candidates per case, selected from five pooled searches, all had higher OASis humanness, predicted Tm, and aggregation scores than their parents (Table S3). Their VH and VL alignments with references appear in Fig. 3F, G for TNF-α and Figs. S3–S6 for the other four cases. Mean OASis scores ranged from 0.647 to 0.731, exceeding parental and Hu-mAb values but remaining below experimental and Sapiens values in every case (Table S3). Selected candidates had higher aggregation scores than all three humanized references in each case. Figure 5 shows component scores of the five highest-reward candidates from run 1 per case.

**Figure 5.**
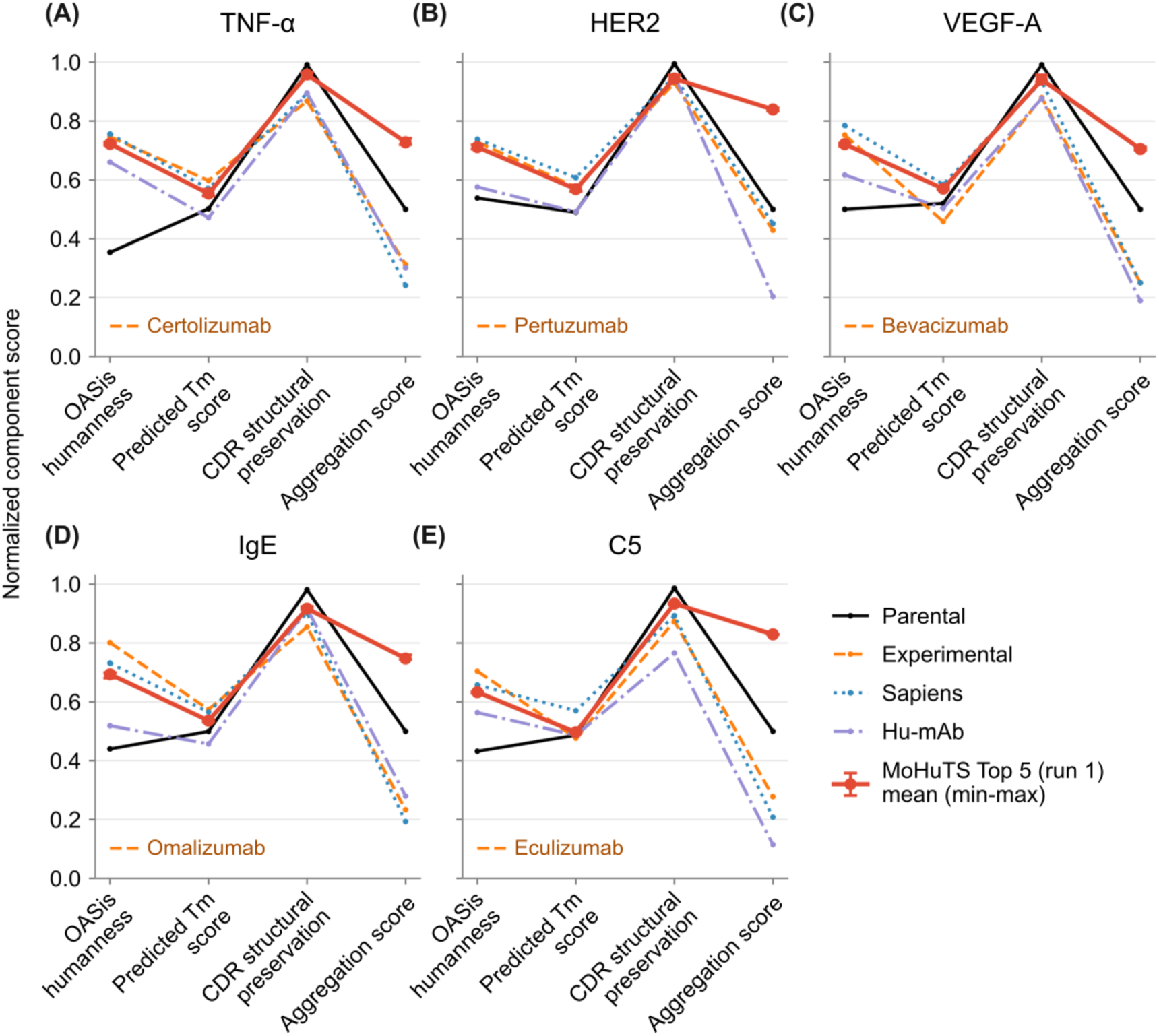
Component scores of the five highest-reward MoHuTS candidates from run 1 and reference sequences. Panels A–E show antibodies targeting TNF-α, HER2, VEGF-A, IgE, and C5, respectively. Red points show means across the five highest-reward MoHuTS candidates from run 1 per case; error bars span their minimum and maximum values. Reference lines represent individual parental, experimentally validated humanized (orange dashed lines labeled by antibody name), Sapiens, and Hu-mAb sequences. Predicted Tm is normalized. Lines connect the four-component scores for each reference or candidate mean.

Sequence substitutions appear in Fig. 3F, G and Figs. S2–S6; Table S3 summarizes total VH/VL counts. The five selected candidates per case had fewer substitutions from parental sequences than experimentally validated humanized, Sapiens-generated, and Hu-mAb-generated references. Mean counts ranged from 18.0 to 22.0, versus 41–60 for experimental references, 29–30 for Sapiens, and 23–40 for Hu-mAb (Table S3). In every case, MoHuTS mean counts were below each corresponding reference count, although substitution count was not included in the reward. Fewer substitutions may reduce unintended effects on parental antibody function (Marks et al., 2021).

Most substitutions also occurred in reference sequences, but their combinations differed (Fig. 3F, G; Figs. S3–S6). Selection using the four-component reward yielded combinations that differed from Sapiens references and included substitutions shared with experimentally validated humanized references. At many positions where Sapiens and experimental references differed, MoHuTS candidates retained parental residues. Thus, MoHuTS may improve humanness and predicted biophysical scores while limiting changes to parental sequences.

To assess high-reward sequence diversity, we compared five sequence-diverse representatives selected by clustering per antibody (Figs. S2–S6). They included substitutions absent from the five highest-reward candidates; most of these substitutions were shared with reference sequences (Figs. S2–S6). Under the same search conditions, C5 representatives showed greater diversity than the other four cases, suggesting more flexible framework substitution patterns compatible with high rewards. This is consistent with the reported retention of high-affinity binding to complement component 5 (C5) after CDR grafting without framework back-mutations (Thomas et al., 1996). The anti-C5 parent’s CDRs may therefore tolerate diverse framework substitutions.

### 3.3 Ablation study

We compared full-reward MoHuTS with single-objective searches optimizing OASis humanness alone (OASis-only). OASis-only searches assigned weights of 1 to humanness and 0 to other components; all other calculation conditions were unchanged.

Full-reward searches achieved higher integrated rewards and lower framework substitution counts than OASis-only searches, whose candidates were retrospectively evaluated with the four-component reward (Fig. 6). Across five cases, rewards averaged across five runs, each summarized over the final 200 candidates, ranged from 0.671 to 0.728 for full-reward searches and 0.530 to 0.639 for OASis-only candidates (Table S1). Full-reward searches yielded 0.89–4.52 fewer framework substitutions and aggregation scores 0.273–0.475 higher, but OASis scores 0.046–0.095 lower. Reward and aggregation scores favored full-reward searches in all 25 matched-seed pairs; substitution counts were lower in 24 of 25 pairs. Predicted *T_m_* and CDR structural preservation differences varied by case. Fig. S7 and Table S1 provide trajectories and mean (SD) summaries. These results suggest that multiple objectives enable more balanced predicted property profiles than humanness alone.

**Figure 6.**
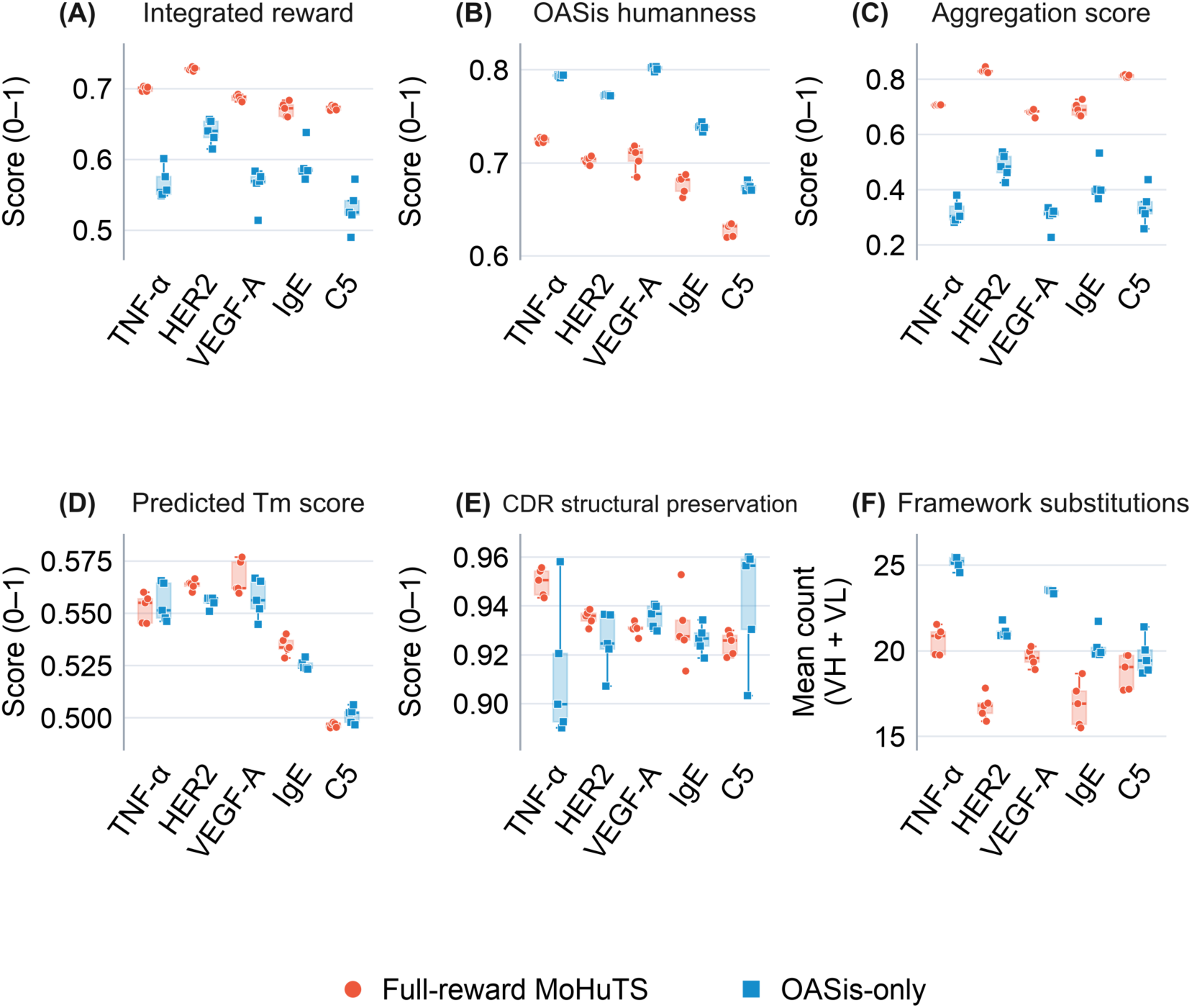
Full-reward and OASis-only searches across five humanization cases (n = 5 per condition and case). Panels A–F show integrated reward, OASis humanness, aggregation score, predicted *T_m_* score, CDR structural preservation score, and total parent-relative VH/VL framework substitutions, respectively. Each symbol is the mean over the final 200 candidates (explored candidates 2,801–3,000) from one independent 3,000-sequence search. Red circles indicate full-reward MoHuTS; blue squares indicate OASis-only optimization. Boxes show the interquartile range and median across five run means; whiskers extend to the most extreme values within 1.5 times the interquartile range.

### 3.4 Additional analyses of antibody–antigen interfaces and sequence humanness

Although low CDR RMSD in predicted structures suggests preserved CDR conformations, it does not guarantee retained antigen binding. To complement this structural assessment, we evaluated modeled antibody–antigen interfaces of the highest-reward candidates using Rosetta metrics previously used to assess antibody–antigen interactions (Adolf-Bryfogle et al., 2018; Schoeder et al., 2021). Changes were case-dependent (Fig. 7). Median interface energies were close to parental values for IgE (−0.43 REU), VEGF-A (−1.93 REU), and C5 (+1.09 REU). TNF-α and HER2 showed positive median differences of +5.48 and +3.27 REU, respectively, indicating higher modeled energies and potentially weaker binding. In the TNF-α and HER2 cases, shape complementarity remained close to parental values despite higher interface energies. For TNF-α, median shape complementarity was 0.687 versus 0.672 for the parent, but median buried interface area was approximately 12% smaller (1,700 versus 1,923 Å²). For HER2, shape complementarity (0.615 versus 0.638) and buried interface area (2,445 versus 2,432 Å²) remained close to parental values (Table S2). Higher interface energies and reduced interface area in some cases warrant experimental evaluation of antigen binding.

**Figure 7.**
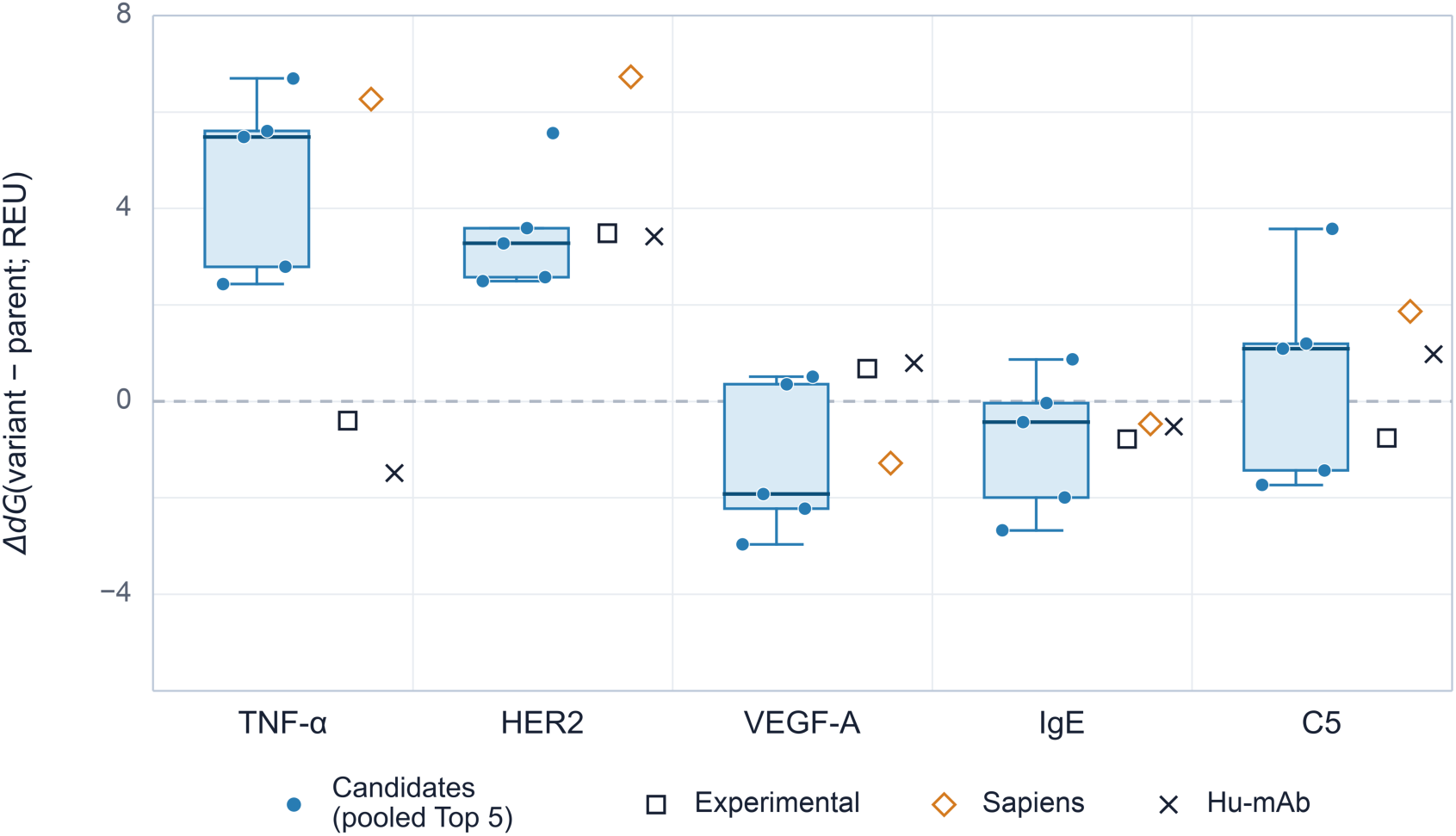
Parent-relative Rosetta antibody–antigen interface scores for the five highest-reward distinct VH/VL sequences selected from five pooled independent searches per case and reference sequences. Each point is a variant’s median dG across three Rosetta runs minus the corresponding three-run parental median. Positive values indicate higher modeled interface energy and potentially weaker binding. Blue points show candidates. Boxes show their interquartile range and median; whiskers extend to the most extreme values within 1.5 times the interquartile range. Open squares, open diamonds, and crosses indicate experimentally validated humanized, Sapiens-generated, and Hu-mAb-generated references, respectively. The dashed line marks the parental baseline (Parent = 0). Negative differences indicate lower modeled interface energy. Values are in Rosetta energy units (REU).

We assessed sequence humanness using two metrics outside the reward: human germline framework identity (Dunbar and Deane, 2016) and AbNatiV scores (Ramon et al., 2024). Germline framework identity was calculated against the closest human V-and joining (J)-gene references over the international ImMunoGeneTics information system (IMGT) framework regions using ANARCI (Dunbar and Deane, 2016). AbNatiV scores used the v1 VH and kappa light-chain variable-domain (VKappa) models (Ramon et al., 2024). Both measures were averaged across VH and VL. Across five searches in all five cases, mean values for the reward top five (five candidates per search; 25 records total) and top 10% (300 candidates per search; 1,500 records total) exceeded parental values (Table 1).

**Table 1.** Humanness measures for generated candidates and references. Germline framework (FR) identity and AbNatiV scores are VH/VL means. “Reward Top 5 per run” averages 25 candidate records: the five highest-reward candidates from each of five searches per antibody (n = 5; 3,000 candidates per search). “Reward Top 10% per run” averages 1,500 records: the 300 highest-reward candidates per search. *OASis humanness was included in the reward.

| **Antibody** | **Metric** | **Parental** | **Reward Top 5 per run** | **Reward Top 10% per run** | **Hu-mAb** | **Sapiens** | **Experimental** |
|---|---|---|---|---|---|---|---|
| Certolizumab | Germline FR identity (%) | 72.2% | 81.8% | 81.3% | 77.3% | 87.2% | 90.6% |
|  | AbNatiV score | 0.633 | 0.802 | 0.791 | 0.745 | 0.879 | 0.828 |
|  | OASis humanness score* | 0.354 | 0.727 | 0.722 | 0.660 | 0.756 | 0.746 |
| Pertuzumab | Germline FR identity (%) | 68.9% | 77.9% | 77.7% | 77.8% | 82.8% | 90.0% |
|  | AbNatiV score | 0.628 | 0.750 | 0.749 | 0.742 | 0.842 | 0.850 |
|  | OASis humanness score* | 0.538 | 0.710 | 0.704 | 0.576 | 0.738 | 0.729 |
| Bevacizumab | Germline FR identity (%) | 76.7% | 84.3% | 83.9% | 80.6% | 91.7% | 91.2% |
|  | AbNatiV score | 0.735 | 0.818 | 0.817 | 0.797 | 0.907 | 0.860 |
|  | OASis humanness score* | 0.500 | 0.716 | 0.701 | 0.617 | 0.785 | 0.752 |
| Omalizumab | Germline FR identity (%) | 72.8% | 80.5% | 80.2% | 82.9% | 88.3% | 92.8% |
|  | AbNatiV score | 0.633 | 0.714 | 0.709 | 0.715 | 0.808 | 0.837 |
|  | OASis humanness score* | 0.440 | 0.686 | 0.674 | 0.519 | 0.731 | 0.801 |
| Eculizumab | Germline FR identity (%) | 70.6% | 79.4% | 79.0% | 84.5% | 85.6% | 92.8% |
|  | AbNatiV score | 0.626 | 0.754 | 0.744 | 0.805 | 0.826 | 0.868 |
|  | OASis humanness score* | 0.432 | 0.640 | 0.628 | 0.563 | 0.657 | 0.704 |

Reward top-five means were comparable to or higher than Hu-mAb values on both measures in most cases. Both top-five and top-10% means remained below experimental and Sapiens values in all five cases. These results support improved sequence humanness relative to parental sequences, as assessed by two metrics not included in the optimization reward. However, lower scores than humanized and Sapiens references do not necessarily imply greater clinical immunogenicity (Ramon et al., 2024). The immunogenicity of MoHuTS candidates requires further evaluation.

## 4. Conclusion

MoHuTS combines Sapiens-proposed framework substitutions with a reward integrating OASis humanness, predicted thermal stability, CDR structural preservation, and aggregation propensity. Across five cases, candidates had higher integrated rewards than parental and humanized references. Selected candidates had fewer substitutions than experimentally validated humanized, Sapiens-generated, and Hu-mAb-generated references. Full-reward searches also yielded fewer framework substitutions on average than OASis-only searches in all five cases. These results support MoHuTS as an in silico approach for prioritizing humanization candidates for experimental validation by considering humanness and developability.

This study has some limitations. First, the selected candidates were evaluated computationally and have not been experimentally validated. Measurements of thermal stability, aggregation, and antigen binding are needed to determine whether predicted improvements are achieved while improving humanness. Second, search outcomes depend on the accuracy and scope of the scoring functions. OASis quantifies sequence humanness but does not directly measure immunogenicity (Prihoda et al., 2022). The Tm predictor was trained on 165 antibody sequences and achieved a cross-validated RMSE of 5.024 °C, but its ability to rank closely related humanization variants remains to be established. CDR RMSD from predicted structures is subject to modeling uncertainty, and small differences among closely related variants may not be reliably resolved (Abanades et al., 2023; Bashour et al., 2024). Low predicted CDR RMSD therefore does not establish preservation of antigen binding. Similarly, AGGRESCAN estimates intrinsic sequence-based aggregation propensity without explicitly accounting for solvent exposure or conformational dynamics (Conchillo-Solé et al., 2007; Zambrano et al., 2015). Its ability to predict experimental aggregation differences among selected variants remains to be established. The reward also excludes expression yield, solubility, viscosity, and chemical stability. Future work could evaluate complementary scoring approaches, including structure-based aggregation prediction with AGGRESCAN3D (Zambrano et al., 2015) and broader developability profiling with the Therapeutic Antibody Profiler (TAP; Raybould et al., 2019). These comparisons would help assess how scoring functions affect candidate prioritization. Finally, the exploration parameter c and reward weights were fixed across cases. These settings have not been established as optimal; other settings may yield different objective trade-offs.

Future work could improve predictive models, add complementary computational evaluations, and incorporate iterative experimental feedback. Expanding training datasets with public and in-house measurements and refining predictors could improve accuracy across diverse antibodies. When proprietary data cannot be shared directly, federated learning may enable collaborative training without centrally pooling datasets (McMahan et al., 2017). Molecular dynamics simulations and structure-based energy evaluations could complement current scoring functions (Choi et al., 2015; Margreitter et al., 2016). Thermal stability, aggregation, and antigen binding measurements could then update predictors and guide subsequent sequence design, creating an iterative cycle of design, testing, and model refinement (Grace et al., 2025).

## Data availability

Antibody sequences and computational results will be deposited at https://github.com/ycu-iil/MoHuTS, including component scores, integrated rewards, and numerical results underlying figures and tables. Published datasets and structural templates are identified in the Methods and Supporting Information.

## Code availability

The MoHuTS source code and configuration files used in this study will be made available at https://github.com/ycu-iil/MoHuTS.

## Acknowledgments

We thank Prof. Yutaka Saito for the helpful discussion. This work was supported by the Japan Science and Technology Agency (JST) Fusion Oriented Research for disruptive Science and Technology (FOREST) Program [grant number JPMJFR232U], the Japan Agency for Medical Research and Development (AMED) “Development of Drug discovery AI platform through Industry-academia Collaboration-DX (DAIIA-X)” [grant number JP26nk0101112], and the Ministry of Education, Culture, Sports, Science and Technology (MEXT) Next-Generation Computational Science Global Reach & Innovation Program Japan [grant number JPMXP19 26254282]. This study was partially supported by the Japan Society for the Promotion of Science Program for Forming Japan’s Peak Research Universities (J-PEAKS) [grant number JPJS00420240015].

## Author contributions

K. Y. and K. T. conceived the study. K. Y. led MoHuTS design and implementation, performed the computational analyses, prepared the figures, and wrote the first draft of the manuscript. S. F. provided technical guidance on implementing the MoHuTS tree search using ChemTSv3. S. I. reviewed the relevant literature and contributed to early discussions of the research plan. K. T. supervised the study, helped interpret the results, and revised the manuscript. All authors reviewed and approved the final version of the manuscript.

