## Supporting Information for "MoHuTS: Developability-Aware Antibody Humanization Using Multi-Objective Tree Search"

### S1 Tm model development and evaluation

We developed the Tm prediction model used in MoHuTS by training a LightGBM regressor (Ke et al., 2017) on AbLang2-paired representations of antibody VH/VL sequences (Olsen et al., 2024). The training dataset comprised 122 clinical-stage antibody records from a pre-existing analysis table containing a subset of the original 137 records in Jain et al. (2017), and 43 human B-cell-derived antibody records from Shehata et al. (2019) with Tm ≤ 65 °C. Target values were Fab Tm measured by differential scanning fluorimetry in the Jain dataset and apparent melting temperature (TmApp) in the Shehata dataset. AbLang2-paired was run in seqcoding mode, with no additional feature scaling at inference. LightGBM used default hyperparameters, including 100 boosting iterations, a learning rate of 0.1, and a maximum of 31 leaves. Five-fold cross-validation used record-level KFold splits with shuffle = True and random_state = 0, and performance was calculated from pooled held-out predictions. This yielded a Pearson correlation coefficient of $r = 0.660$ and a root mean squared error (RMSE) of 5.024 °C. The final model used in MoHuTS was trained on the full dataset.

**Table S1. Final-window comparison of full-reward and OASis-only searches across five antibody humanization cases (n = 5 per condition).**

| Case | Search | Reward | OASis | Tm | CDR | Agg. | Subst. |
|---|---|---|---|---|---|---|---|
| Certolizumab | Full reward | 0.700 (0.003) | 0.725 (0.003) | 0.553 (0.007) | 0.950 (0.006) | 0.706 (0.001) | 20.61 (0.80) |
| Certolizumab | OASis only | 0.567 (0.022) | 0.794 (0.001) | 0.555 (0.009) | 0.912 (0.028) | 0.319 (0.041) | 25.13 (0.38) |
| Pertuzumab | Full reward | 0.728 (0.002) | 0.703 (0.004) | 0.564 (0.002) | 0.935 (0.003) | 0.831 (0.008) | 16.76 (0.72) |
| Pertuzumab | OASis only | 0.639 (0.017) | 0.773 (0.001) | 0.556 (0.003) | 0.925 (0.012) | 0.485 (0.044) | 21.13 (0.38) |
| Bevacizumab | Full reward | 0.687 (0.004) | 0.706 (0.013) | 0.567 (0.008) | 0.931 (0.003) | 0.681 (0.012) | 19.60 (0.53) |
| Bevacizumab | OASis only | 0.562 (0.028) | 0.801 (0.002) | 0.557 (0.009) | 0.936 (0.005) | 0.301 (0.043) | 23.48 (0.09) |
| Omalizumab | Full reward | 0.671 (0.010) | 0.677 (0.010) | 0.535 (0.004) | 0.931 (0.014) | 0.692 (0.025) | 16.89 (1.32) |
| Omalizumab | OASis only | 0.593 (0.026) | 0.739 (0.004) | 0.525 (0.002) | 0.926 (0.006) | 0.419 (0.065) | 20.28 (0.82) |
| Eculizumab | Full reward | 0.673 (0.003) | 0.628 (0.007) | 0.497 (0.001) | 0.925 (0.005) | 0.812 (0.005) | 18.79 (1.01) |
| Eculizumab | OASis only | 0.530 (0.030) | 0.675 (0.005) | 0.501 (0.004) | 0.942 (0.025) | 0.338 (0.065) | 19.69 (1.09) |

Note. Values are means of five independent run means (sample SD in parentheses). Each run mean uses evaluations 2,801–3,000 from a 3,000-pair search; runs are replicates, with seeds matched between conditions. Full integrated rewards for OASis-only candidates were calculated retrospectively. Substitution counts sum parent-relative VH and VL changes. Protected Kabat CDR residues were unchanged in all 150,000 candidate records across 50 runs; pairs may recur across runs. Agg. and Subst. denote aggregation score and framework substitution count; Tm and CDR denote normalized predicted melting-temperature and CDR preservation scores. Displayed values are rounded; text comparisons use unrounded run means.

### Table S2. Rosetta interface scores for selected candidates and reference sequences

| Antibody | PDB code | Heavy-chain ID | Light-chain ID | Antigen chains | Interface |
|---|---|---|---|---|---|
| Certolizumab | 5WUX | A | B | F | AB_F |
| Pertuzumab | 1S78 | D | C | A | CD_A |
| Omalizumab | 5HYS | A | B | G | AB_G |
| Eculizumab | 5I5K | H | L | B | B_HL |
| Bevacizumab | 1BJ1 | H | L | V/W | HL_VW |

| Antibody | Sequence | dG Run 1 (REU) | dG Run 2 (REU) | dG Run 3 (REU) | ΔdG (REU) | Median dSASA_int (Å²) | Median SC |
|---|---|---|---|---|---|---|---|
| **Certolizumab** | Parental | -59.190 | -58.637 | -62.138 | 0.000 | 1,922.7 | 0.672 |
| | Experimental | -63.099 | -59.593 | -53.754 | -0.403 | 1,862.4 | 0.671 |
| | Sapiens | -57.217 | -52.785 | -52.925 | +6.265 | 1,719.3 | 0.679 |
| | Hu-mAb | -60.481 | -62.322 | -60.680 | -1.490 | 1,944.7 | 0.687 |
| | MoHuTS rank 1 | -53.587 | -57.336 | -53.133 | +5.603 | 1,759.5 | 0.631 |
| | MoHuTS rank 2 | -53.710 | -56.398 | -51.635 | +5.480 | 1,695.1 | 0.687 |
| | MoHuTS rank 3 | -56.400 | -53.732 | -57.347 | +2.790 | 1,690.9 | 0.703 |
| | MoHuTS rank 4 | -60.739 | -55.786 | -56.761 | +2.429 | 1,724.5 | 0.693 |
| | MoHuTS rank 5 | -52.498 | -52.246 | -53.774 | +6.692 | 1,700.4 | 0.660 |
| **Pertuzumab** | Parental | -57.838 | -57.124 | -53.866 | 0.000 | 2,432.2 | 0.638 |
| | Experimental | -53.641 | -51.599 | -56.556 | +3.483 | 2,526.1 | 0.632 |
| | Sapiens | -52.707 | -49.995 | -50.396 | +6.728 | 2,290.9 | 0.584 |
| | Hu-mAb | -54.816 | -53.708 | -52.948 | +3.416 | 2,484.7 | 0.587 |
| | MoHuTS rank 1 | -55.662 | -52.226 | -53.535 | +3.589 | 2,497.9 | 0.615 |
| | MoHuTS rank 2 | -53.487 | -53.851 | -58.813 | +3.273 | 2,448.7 | 0.562 |
| | MoHuTS rank 3 | -54.633 | -50.591 | -85.136 | +2.491 | 2,433.0 | 0.578 |
| | MoHuTS rank 4 | -54.551 | -48.192 | -57.809 | +2.573 | 2,406.4 | 0.627 |
| | MoHuTS rank 5 | -48.481 | -51.564 | -52.045 | +5.560 | 2,445.4 | 0.625 |

| Antibody | Sequence | dG Run 1 (REU) | dG Run 2 (REU) | dG Run 3 (REU) | ΔdG (REU) | Median dSASA_int (Å²) | Median SC |
|---|---|---|---|---|---|---|---|
| **Omalizumab** | Parental | -37.080 | -35.585 | -37.735 | 0.000 | 1,151.3 | 0.706 |
| | Experimental | -37.864 | -35.874 | -39.167 | -0.784 | 1,217.3 | 0.710 |
| | Sapiens | -37.794 | -37.550 | -36.634 | -0.470 | 1,162.8 | 0.701 |
| | Hu-mAb | -37.904 | -36.608 | -37.606 | -0.526 | 1,148.2 | 0.687 |
| | MoHuTS rank 1 | -37.640 | -37.513 | -35.805 | -0.433 | 1,076.3 | 0.693 |
| | MoHuTS rank 2 | -36.587 | -39.122 | -37.118 | -0.038 | 1,136.9 | 0.730 |
| | MoHuTS rank 3 | -36.822 | -39.757 | -40.274 | -2.677 | 1,170.0 | 0.722 |
| | MoHuTS rank 4 | -38.319 | -36.211 | -35.457 | +0.869 | 1,072.1 | 0.726 |
| | MoHuTS rank 5 | -39.076 | -36.581 | -40.653 | -1.996 | 1,156.7 | 0.727 |
| **Eculizumab** | Parental | -39.654 | -39.423 | -40.426 | 0.000 | 1,840.0 | 0.585 |
| | Experimental | -40.416 | -42.252 | -36.186 | -0.762 | 1,847.5 | 0.638 |
| | Sapiens | -37.338 | -42.804 | -37.791 | +1.863 | 1,819.7 | 0.643 |
| | Hu-mAb | -40.779 | -38.679 | -36.464 | +0.975 | 1,888.2 | 0.680 |
| | MoHuTS rank 1 | -38.864 | -41.389 | -41.813 | -1.735 | 1,918.4 | 0.615 |
| | MoHuTS rank 2 | -26.545 | -40.722 | -36.079 | +3.575 | 1,805.7 | 0.585 |
| | MoHuTS rank 3 | -38.564 | -37.272 | -39.568 | +1.090 | 1,809.2 | 0.648 |
| | MoHuTS rank 4 | -41.220 | -40.799 | -41.090 | -1.436 | 1,860.3 | 0.605 |
| | MoHuTS rank 5 | -36.899 | -41.095 | -38.459 | +1.195 | 1,804.1 | 0.607 |
| **Bevacizumab** | Parental | -58.366 | -55.414 | -54.594 | 0.000 | 1,675.8 | 0.769 |
| | Experimental | -51.359 | -60.866 | -54.735 | +0.679 | 1,608.5 | 0.773 |
| | Sapiens | -59.227 | -56.698 | -55.296 | -1.284 | 1,643.8 | 0.791 |
| | Hu-mAb | -58.822 | -54.623 | -53.224 | +0.791 | 1,604.4 | 0.789 |
| | MoHuTS rank 1 | -50.216 | -55.459 | -54.906 | +0.508 | 1,643.1 | 0.743 |
| | MoHuTS rank 2 | -57.641 | -61.346 | -55.312 | -2.227 | 1,667.3 | 0.792 |
| | MoHuTS rank 3 | -56.797 | -52.565 | -55.064 | +0.350 | 1,749.4 | 0.792 |
| | MoHuTS rank 4 | -58.382 | -58.701 | -54.819 | -2.968 | 1,737.9 | 0.801 |
| | MoHuTS rank 5 | -61.877 | -57.339 | -49.147 | -1.925 | 1,778.7 | 0.739 |

Note. ΔdG is each sequence's three-run median dG minus the three-run parental median. The final two columns show three-run medians of buried interface area (dSASA_int, Å²) and shape complementarity (SC; sc_value, dimensionless), respectively. Energy values are in REU.

**Table S3. Integrated rewards, component scores, and VH/VL substitution counts for MoHuTS candidates and reference sequences**

| Antibody | Sequence | Reward | OASis | Tm | CDR score | Agg. | Total subst. | CDR subst. |
|---|---|---|---|---|---|---|---|---|
| Certolizumab | Parental | 0.4667 | 0.3541 | 0.5018 | 0.9912 | 0.5000 | 0 | 0 |
| | Experimental | 0.5580 | 0.7464 | 0.5978 | 0.8678 | 0.3120 | 51 | 3 |
| | Sapiens | 0.5163 | 0.7560 | 0.5708 | 0.8937 | 0.2418 | 29 | 0 |
| | Hu-mAb | 0.5027 | 0.6603 | 0.4719 | 0.8955 | 0.3007 | 28 | 4 |
| | MoHuTS Top1 | 0.7147 | 0.7368 | 0.5730 | 0.9644 | 0.7197 | 22 | 0 |
| | MoHuTS Top2 | 0.7146 | 0.7273 | 0.5683 | 0.9516 | 0.7390 | 22 | 0 |
| | MoHuTS Top3 | 0.7143 | 0.7321 | 0.5700 | 0.9641 | 0.7271 | 22 | 0 |
| | MoHuTS Top4 | 0.7140 | 0.7321 | 0.5677 | 0.9494 | 0.7318 | 22 | 0 |
| | MoHuTS Top5 | 0.7140 | 0.7273 | 0.5732 | 0.9647 | 0.7294 | 22 | 0 |
| | **MoHuTS Top5 mean** | **0.7143** | **0.7311** | **0.5704** | **0.9588** | **0.7294** | **22.0** | **0.0** |
| Pertuzumab | Parental | 0.5493 | 0.5381 | 0.4899 | 0.9943 | 0.5000 | 0 | 0 |
| | Experimental | 0.6074 | 0.7286 | 0.5750 | 0.9292 | 0.4290 | 52 | 0 |
| | Sapiens | 0.6285 | 0.7381 | 0.6077 | 0.9523 | 0.4516 | 29 | 0 |
| | Hu-mAb | 0.4295 | 0.5762 | 0.4910 | 0.9525 | 0.2036 | 29 | 4 |
| | MoHuTS Top1 | 0.7414 | 0.7048 | 0.5733 | 0.9372 | 0.8706 | 18 | 0 |
| | MoHuTS Top2 | 0.7407 | 0.7190 | 0.5664 | 0.9409 | 0.8510 | 18 | 0 |
| | MoHuTS Top3 | 0.7404 | 0.7095 | 0.5642 | 0.9358 | 0.8687 | 18 | 0 |
| | MoHuTS Top4 | 0.7402 | 0.7048 | 0.5603 | 0.9418 | 0.8781 | 18 | 0 |
| | MoHuTS Top5 | 0.7402 | 0.7095 | 0.5624 | 0.9331 | 0.8706 | 18 | 0 |
| | **MoHuTS Top5 mean** | **0.7406** | **0.7095** | **0.5653** | **0.9378** | **0.8678** | **18.0** | **0.0** |
| Bevacizumab | Parental | 0.5396 | 0.5000 | 0.5200 | 0.9910 | 0.5000 | 0 | 0 |
| | Experimental | 0.4978 | 0.7523 | 0.4587 | 0.8798 | 0.2507 | 41 | 0 |
| | Sapiens | 0.5347 | 0.7850 | 0.5847 | 0.9344 | 0.2506 | 29 | 0 |
| | Hu-mAb | 0.4305 | 0.6168 | 0.5035 | 0.8768 | 0.1894 | 23 | 3 |
| | MoHuTS Top1 | 0.7056 | 0.7243 | 0.5833 | 0.9465 | 0.7014 | 21 | 0 |
| | MoHuTS Top2 | 0.7037 | 0.7196 | 0.5677 | 0.9589 | 0.7111 | 21 | 0 |
| | MoHuTS Top3 | 0.7036 | 0.7290 | 0.5641 | 0.9380 | 0.7067 | 21 | 0 |
| | MoHuTS Top4 | 0.7033 | 0.7243 | 0.5773 | 0.9360 | 0.7014 | 21 | 0 |
| | MoHuTS Top5 | 0.7032 | 0.7196 | 0.5668 | 0.9548 | 0.7111 | 21 | 0 |
| | **MoHuTS Top5 mean** | **0.7039** | **0.7234** | **0.5719** | **0.9468** | **0.7063** | **21.0** | **0.0** |

**Table S3 (continued). Integrated rewards, component scores, and VH/VL substitution counts**

| Antibody | Sequence | Reward | OASis | Tm | CDR score | Agg. | Total subst. | CDR subst. |
|---|---|---|---|---|---|---|---|---|
| Omalizumab | Parental | 0.5080 | 0.4398 | 0.4997 | 0.9807 | 0.5000 | 0 | 0 |
| | Experimental | 0.5211 | 0.8009 | 0.5743 | 0.8542 | 0.2335 | 60 | 7 |
| | Sapiens | 0.4756 | 0.7315 | 0.5652 | 0.9018 | 0.1929 | 30 | 0 |
| | Hu-mAb | 0.4450 | 0.5185 | 0.4571 | 0.9171 | 0.2802 | 40 | 6 |
| | MoHuTS Top1 | 0.6965 | 0.7037 | 0.5403 | 0.9119 | 0.7438 | 19 | 0 |
| | MoHuTS Top2 | 0.6959 | 0.7037 | 0.5406 | 0.9036 | 0.7438 | 19 | 0 |
| | MoHuTS Top3 | 0.6953 | 0.6991 | 0.5407 | 0.9183 | 0.7438 | 19 | 0 |
| | MoHuTS Top4 | 0.6947 | 0.6991 | 0.5409 | 0.9103 | 0.7438 | 19 | 0 |
| | MoHuTS Top5 | 0.6946 | 0.6806 | 0.5402 | 0.9140 | 0.7704 | 19 | 0 |
| | **MoHuTS Top5 mean** | **0.6954** | **0.6972** | **0.5405** | **0.9116** | **0.7492** | **19.0** | **0.0** |
| Eculizumab | Parental | 0.5020 | 0.4319 | 0.4868 | 0.9858 | 0.5000 | 0 | 0 |
| | Experimental | 0.5036 | 0.7042 | 0.4768 | 0.8734 | 0.2780 | 43 | 0 |
| | Sapiens | 0.4663 | 0.6573 | 0.5698 | 0.8926 | 0.2080 | 29 | 0 |
| | Hu-mAb | 0.3501 | 0.5634 | 0.4864 | 0.7654 | 0.1149 | 27 | 4 |
| | MoHuTS Top1 | 0.6883 | 0.6526 | 0.4981 | 0.9305 | 0.8292 | 20 | 0 |
| | MoHuTS Top2 | 0.6863 | 0.6479 | 0.4982 | 0.9296 | 0.8292 | 20 | 0 |
| | MoHuTS Top3 | 0.6863 | 0.6479 | 0.4973 | 0.9331 | 0.8292 | 20 | 0 |
| | MoHuTS Top4 | 0.6863 | 0.6479 | 0.5042 | 0.9407 | 0.8191 | 20 | 0 |
| | MoHuTS Top5 | 0.6858 | 0.6385 | 0.5146 | 0.9275 | 0.8262 | 20 | 0 |
| | **MoHuTS Top5 mean** | **0.6866** | **0.6469** | **0.5025** | **0.9323** | **0.8266** | **20.0** | **0.0** |

Note. Top1–Top5 are the five highest-reward distinct VH/VL pairs per antibody, selected by pooling five independent full-reward searches (3,000 unique pairs per run), deduplicating exact pairs, and ranking by median reward across runs containing each pair. Each selected pair occurred in one run. These candidates correspond to Figs. 3F,G and S2–S6. MoHuTS Top5 mean averages these five pairs in each column using unrounded values; it differs from the per-run Top5 summary in main-text Table 1. References are individual sequences; Experimental denotes the experimentally validated humanized reference. Reward denotes integrated reward; OASis, Tm, CDR score, and Agg. denote humanness, normalized predicted Tm, CDR preservation, and aggregation scores, respectively. Scores range from 0 to 1; higher is more favorable. Tm is normalized, not in °C. Total subst. and CDR subst. count parent-relative amino acid substitutions across VH/VL and within Kabat CDRs, respectively. Selected MoHuTS candidates have zero CDR substitutions. Scores and mean substitution counts are rounded to four and one decimal places, respectively.

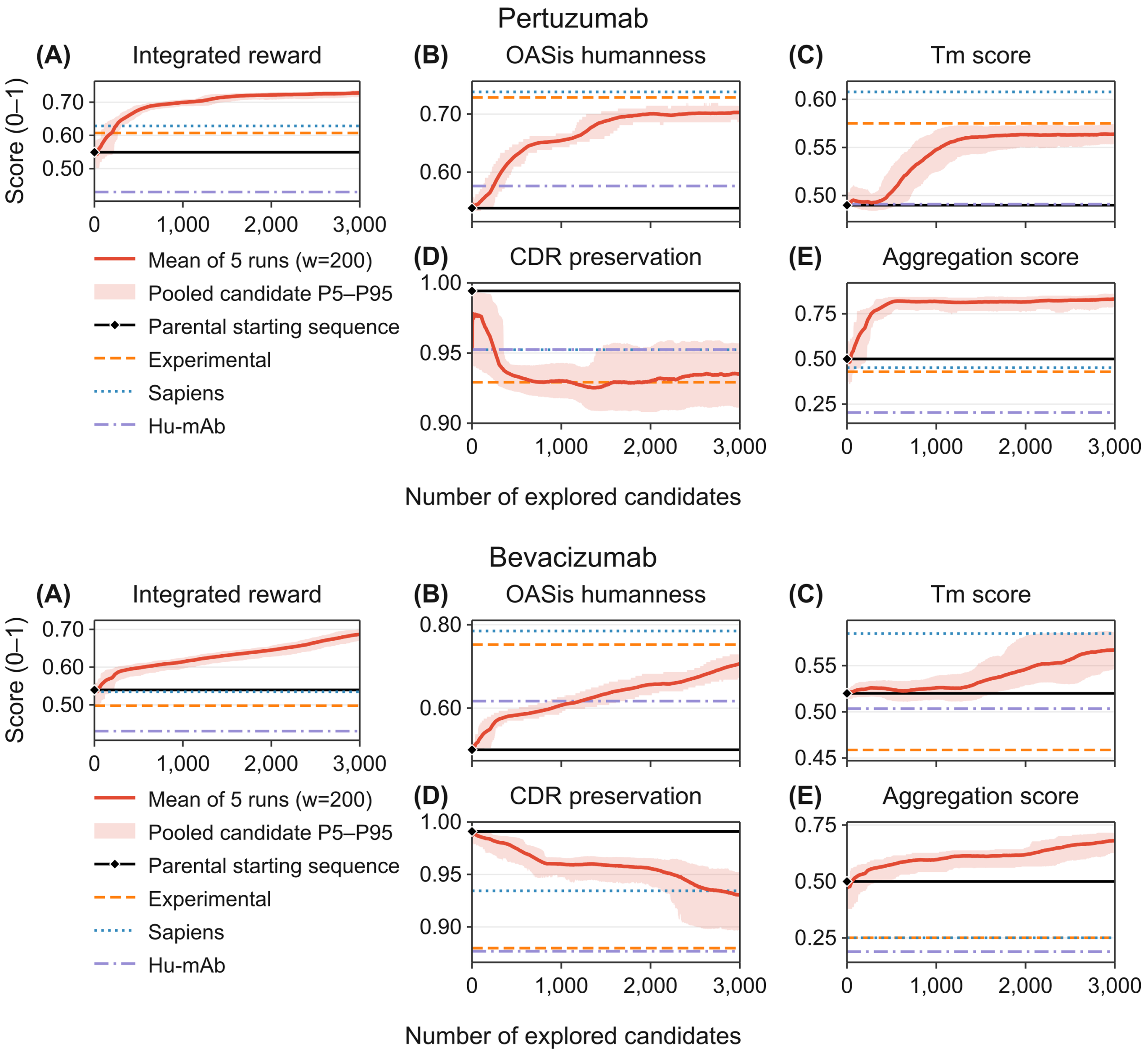


**Figure S1. Integrated reward and component-score trajectories for four antibody cases.** For each antibody, panels A–E show integrated reward, OASis humanness, normalized predicted Tm, CDR preservation, and aggregation scores, respectively. Solid red lines show trailing 200-candidate means averaged across five independent runs; means for the first 199 candidates use all candidates explored so far per run. Shading shows the 5th–95th percentiles pooled across runs within these windows, reflecting candidate variation, not confidence intervals across runs. The x-axis shows unique candidates explored per run (3,000 per run). Black diamonds mark parental scores at x = 0. Horizontal lines indicate parental, experimentally validated humanized (Experimental), Sapiens, and Hu-mAb references. Higher scores are more favorable.

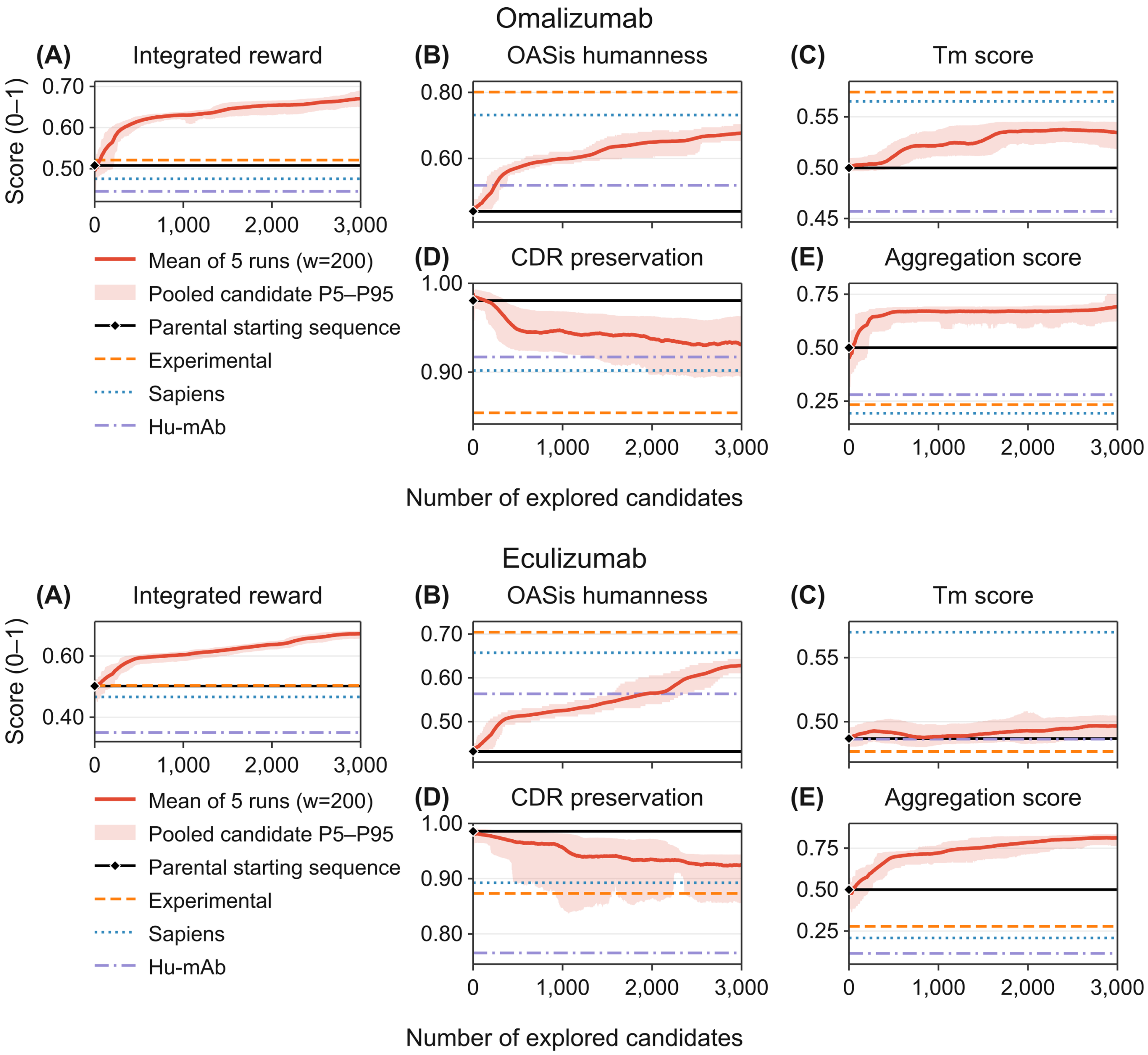


**Figure S1 (continued).** Omalizumab and eculizumab trajectories.

**(A)** Certolizumab · VH

| Candidate / Method | Reward | Rank | Mutations | H1 | H2 | H6 | H9 | H10 | H12 | H13 | H16 | H17 | H18 | H19 | H20 | H23 | H28 | H38 | H43 | H44 | H45 | H46 | H48 | H49 | H60* | H62* | H63* | H68 | H71 | H72 | H75 | H76 | H78 | H79 | H82 | H82B | H83 | H84 | H85 | H89 | H91 | H108 |
|---|---|---|---|---|---|---|---|---|---|---|---|---|---|---|---|---|---|---|---|---|---|---|---|---|---|---|---|---|---|---|---|---|---|---|---|---|---|---|---|---|---|---|
| **Parental** | | | 0 | Q | I | Q | P | E | K | K | E | T | V | K | I | K | V | K | K | A | F | K | M | G | V | D | F | A | L | E | A | S | A | F | I | N | K | N | E | T | F | S |
| Top1 | 0.7147 | #1 | 9 | · | · | · | · | · | · | · | · | S | · | · | · | · | T | R | · | G | · | E | · | · | · | · | · | · | · | D | · | N | · | Y | · | · | · | · | · | · | Y | · |
| Top2 | 0.7146 | #2 | 11 | · | · | · | · | · | · | · | · | S | · | · | V | · | T | R | · | G | L | E | · | · | · | · | · | · | · | D | · | N | · | Y | · | · | · | · | · | · | Y | · |
| Top3 | 0.7143 | #3 | 10 | · | · | · | · | · | · | · | · | S | · | · | V | · | T | R | · | G | · | E | · | · | · | · | · | · | · | D | · | N | · | Y | · | · | · | · | · | · | Y | · |
| Top4 | 0.7140 | #4 | 10 | · | · | · | · | · | · | · | · | S | · | · | · | · | T | R | · | G | L | E | · | · | · | · | · | · | · | D | · | N | · | Y | · | · | · | · | · | · | Y | · |
| Top5 | 0.7140 | #5 | 10 | · | · | · | · | · | · | · | · | S | · | · | · | · | T | R | Q | G | · | E | · | · | · | · | · | · | · | D | · | N | · | Y | · | · | · | · | · | · | Y | · |
| D1 | 0.7147 | #1 | 9 | · | · | · | · | · | · | · | · | S | · | · | · | · | T | R | · | G | · | E | · | · | · | · | · | · | · | D | · | N | · | Y | · | · | · | · | · | · | Y | · |
| D2 | 0.7130 | #12 | 11 | · | V | · | · | · | · | · | · | S | · | · | · | · | T | R | · | G | L | E | · | · | · | · | · | · | · | D | · | N | · | Y | · | · | · | · | · | · | Y | · |
| D3 | 0.7098 | #113 | 11 | · | · | · | · | · | · | · | · | S | · | · | · | · | T | R | · | G | L | E | · | · | · | · | · | · | · | D | · | N | · | Y | · | · | Q | · | · | · | Y | · |
| D4 | 0.7029 | #624 | 10 | · | · | · | · | · | · | · | · | S | · | · | V | · | T | R | · | G | · | E | · | · | · | · | · | · | · | D | · | N | · | Y | · | · | · | · | · | · | Y | · |
| D5 | 0.7010 | #964 | 11 | · | · | · | · | · | · | · | · | S | · | · | · | · | T | R | Q | G | · | E | · | · | · | · | · | · | · | D | · | N | · | Y | · | · | · | · | D | · | Y | · |
| Experimental | 0.5580 | — | 31 | E | V | E | G | G | V | Q | G | S | L | R | L | A | · | R | · | G | L | E | · | · | A | S | V | T | · | D | K | · | · | Y | M | S | R | A | · | V | Y | L |
| Sapiens | 0.5163 | — | 14 | · | V | · | S | · | · | · | A | S | · | · | · | · | T | R | · | G | L | E | · | · | · | · | · | V | · | D | · | · | · | · | · | · | · | A | · | V | · | L |
| Hu-mAb | 0.5027 | — | 18 | · | · | · | G | D | · | Q | · | Y | · | R | · | A | · | R | · | · | · | · | L | S | · | S | V | · | R | · | K | N | L | · | · | · | · | A | · | V | · | L |

**(B)** Certolizumab · VL

| Candidate / Method | Reward | Rank | Mutations | L3 | L8 | L9 | L10 | L11 | L13 | L20 | L21 | L24* | L42 | L43 | L46 | L55* | L60 | L63 | L77 | L78 | L80 | L83 | L85 | L87 | L100 | L103 | L104 | L106 |
|---|---|---|---|---|---|---|---|---|---|---|---|---|---|---|---|---|---|---|---|---|---|---|---|---|---|---|---|---|
| **Parental** | | | 0 | V | Q | K | F | M | T | S | V | K | Q | S | A | Y | Y | T | T | V | S | L | E | F | A | K | L | L |
| Top1 | 0.7147 | #1 | 13 | · | P | S | · | L | A | T | I | · | · | · | L | · | S | S | · | L | · | F | T | · | Q | · | · | · |
| Top2 | 0.7146 | #2 | 11 | · | P | S | · | L | A | · | I | · | · | · | L | · | S | · | · | L | · | F | T | · | Q | · | · | · |
| Top3 | 0.7143 | #3 | 12 | · | P | S | · | L | A | T | I | · | · | · | L | · | S | · | · | L | · | F | T | · | Q | · | · | · |
| Top4 | 0.7140 | #4 | 12 | · | P | S | · | L | A | · | I | · | · | · | L | · | S | S | · | L | · | F | T | · | Q | · | · | · |
| Top5 | 0.7140 | #5 | 12 | · | P | S | · | L | A | · | I | · | · | · | L | · | S | · | · | L | · | F | T | · | Q | · | V | · |
| D1 | 0.7147 | #1 | 13 | · | P | S | · | L | A | T | I | · | · | · | L | · | S | S | · | L | · | F | T | · | Q | · | · | · |
| D2 | 0.7130 | #12 | 11 | · | P | S | · | L | A | · | I | · | · | · | L | · | · | · | · | L | · | F | T | · | Q | · | V | · |
| D3 | 0.7098 | #113 | 11 | Q | P | S | · | · | A | · | I | · | · | · | L | · | S | · | · | L | · | F | T | · | Q | · | · | · |
| D4 | 0.7029 | #624 | 10 | · | P | S | · | · | A | · | I | · | · | · | L | · | · | · | S | · | · | F | T | · | Q | R | · | · |
| D5 | 0.7010 | #964 | 8 | Q | · | S | · | · | A | · | I | · | · | · | L | · | · | · | · | · | · | F | T | · | Q | · | · | · |
| Experimental | 0.5580 | — | 20 | Q | P | S | S | L | A | T | I | · | K | A | · | · | · | S | S | L | P | F | T | Y | Q | · | V | I |
| Sapiens | 0.5163 | — | 15 | Q | P | S | · | L | A | T | I | · | · | A | L | · | S | S | · | L | · | · | T | · | Q | · | · | I |
| Hu-mAb | 0.5027 | — | 10 | Q | · | S | · | · | A | · | · | R | · | A | · | Q | · | · | · | L | P | · | T | · | Q | · | · | · |

Aliphatic AVILM | Aromatic FWY | Polar STNQC | Basic KRH | Acidic DE | Gly/Pro GP

**Figure S2. Sequence substitutions in certolizumab candidates and reference sequences.** Panels A and B show VH and VL alignments, respectively. Top1–Top5 are the highest-reward unique VH/VL pairs pooled from five independent searches. Greedy farthest-first sampling selected D1–D5 from pairs with median observed reward ≥98% of the case maximum. Reward is the median across runs containing each pair; ranks refer to pooled unique pairs. Experimental, Sapiens, and Hu-mAb comparators are unranked. Mutation counts are parent-relative substitutions per chain. Columns include all mutated Kabat positions across candidates and comparators. Dots indicate parental residues, colors indicate replacement-residue classes, and asterisks mark Kabat CDR positions.

**(A)** Pertuzumab · VH

| Candidate / Method | Reward | Rank | Mutations | H1 | H5 | H6 | H9 | H10 | H11 | H12 | H13 | H16 | H18 | H19 | H20 | H23 | H27 | H38 | H40 | H41 | H43 | H44 | H48 | H49 | H60* | H63* | H66 | H67 | H68 | H69 | H70 | H71 | H73 | H75 | H76 | H77 | H78 | H80 | H81 | H82 | H82A | H83 | H84 | H85 | H108 | H109 |
|---|---|---|---|---|---|---|---|---|---|---|---|---|---|---|---|---|---|---|---|---|---|---|---|---|---|---|---|---|---|---|---|---|---|---|---|---|---|---|---|---|---|---|---|---|---|---|
| **Parental** | | | 0 | E | Q | Q | P | E | L | V | K | T | V | K | I | K | F | K | S | H | K | S | I | G | N | F | K | A | S | L | T | V | R | S | R | I | V | M | E | L | R | T | F | E | T | L |
| Top1 | 0.7414 | #1 | 12 | · | · | · | · | · | · | K | · | · | · | · | · | · | Y | R | A | · | Q | G | M | · | · | · | · | · | · | M | · | R | · | T | · | T | · | · | · | · | · | · | S | · | · | · |
| Top2 | 0.7407 | #2 | 12 | Q | · | · | · | · | · | K | · | · | · | · | · | · | Y | R | A | · | Q | G | · | · | · | · | · | · | · | M | · | R | · | T | · | T | · | · | · | · | · | · | S | · | · | · |
| Top3 | 0.7404 | #3 | 13 | Q | · | · | · | · | · | K | · | · | · | · | · | · | Y | R | A | · | Q | G | M | · | · | · | · | · | · | M | · | R | · | T | · | T | · | · | · | · | · | · | S | · | · | · |
| Top4 | 0.7402 | #4 | 13 | · | · | · | · | · | · | K | · | · | · | · | · | · | Y | R | A | · | Q | G | M | · | · | · | · | · | · | M | · | R | · | T | · | T | · | · | · | · | · | · | S | D | · | · |
| Top5 | 0.7402 | #5 | 12 | · | · | · | · | · | · | K | · | · | · | · | · | · | Y | R | A | · | Q | G | M | · | · | · | · | · | · | M | · | R | · | T | · | T | · | · | · | · | · | · | S | · | · | · |
| D1 | 0.7414 | #1 | 12 | · | · | · | · | · | · | K | · | · | · | · | · | · | Y | R | A | · | Q | G | M | · | · | · | · | · | · | M | · | R | · | T | · | T | · | · | · | · | · | · | S | · | · | · |
| D2 | 0.7326 | #304 | 10 | Q | · | · | · | · | V | K | · | · | · | · | · | · | · | R | A | · | · | G | · | · | · | · | · | · | · | · | · | R | · | T | · | T | · | · | · | · | · | · | S | · | · | · |
| D3 | 0.7313 | #553 | 10 | · | · | · | · | · | · | K | · | · | · | · | · | · | · | R | A | · | Q | G | · | · | · | · | R | · | · | · | · | R | · | T | · | T | · | · | · | · | · | · | S | · | · | · |
| D4 | 0.7310 | #615 | 12 | · | · | · | · | · | V | K | · | · | · | · | · | · | · | R | A | · | · | G | M | · | · | · | R | · | · | · | · | R | · | T | · | T | · | · | · | · | · | · | S | · | · | V |
| D5 | 0.7269 | #1902 | 12 | Q | · | · | · | · | V | K | · | A | · | · | · | · | · | R | A | · | · | G | M | · | · | · | · | · | · | · | · | R | · | T | · | T | · | · | · | · | · | · | S | · | · | · |
| Experimental | 0.6074 | — | 32 | · | V | E | G | G | · | · | Q | G | L | R | L | A | · | R | A | P | · | G | V | A | · | · | R | F | T | · | S | · | · | K | N | T | L | L | Q | M | N | R | A | · | L | V |
| Sapiens | 0.6285 | — | 14 | · | V | · | · | · | · | K | · | A | · | · | · | · | · | R | A | P | · | G | · | · | · | · | R | F | · | · | · | R | T | · | · | T | · | · | · | · | · | · | · | · | L | V |
| Hu-mAb | 0.4295 | — | 19 | · | L | · | G | T | · | · | · | G | L | R | · | V | · | R | A | · | · | · | V | S | A | G | R | F | · | · | S | · | · | K | · | · | · | · | · | M | · | · | · | · | · | V |

**(B)** Pertuzumab · VL

| Candidate / Method | Reward | Rank | Mutations | L2 | L3 | L8 | L9 | L10 | L11 | L13 | L20 | L24* | L39 | L42 | L43 | L54* | L60 | L63 | L73 | L78 | L80 | L83 | L85 | L100 | L104 |
|---|---|---|---|---|---|---|---|---|---|---|---|---|---|---|---|---|---|---|---|---|---|---|---|---|---|
| **Parental** | | | 0 | T | V | H | K | I | M | T | S | K | R | Q | S | R | D | T | F | V | A | L | V | G | L |
| Top1 | 0.7414 | #1 | 6 | · | Q | P | S | · | · | A | · | · | · | · | · | · | · | · | · | · | · | F | · | Q | · |
| Top2 | 0.7407 | #2 | 6 | · | Q | P | S | · | · | A | · | · | · | · | · | · | · | S | · | · | · | · | · | Q | · |
| Top3 | 0.7404 | #3 | 5 | · | Q | P | S | · | · | A | · | · | · | · | · | · | · | · | · | · | · | · | · | Q | · |
| Top4 | 0.7402 | #4 | 5 | · | Q | P | S | · | · | A | · | · | · | · | · | · | · | · | · | · | · | · | · | Q | · |
| Top5 | 0.7402 | #5 | 6 | · | Q | P | S | · | · | A | · | · | · | · | · | · | · | S | · | · | · | · | · | Q | · |
| D1 | 0.7414 | #1 | 6 | · | Q | P | S | · | · | A | · | · | · | · | · | · | · | · | · | · | · | F | · | Q | · |
| D2 | 0.7326 | #304 | 7 | · | Q | P | S | · | · | A | T | · | · | · | · | · | · | S | · | · | · | · | · | Q | · |
| D3 | 0.7313 | #553 | 7 | · | Q | P | S | · | · | A | · | · | · | · | · | · | · | · | L | L | · | · | · | Q | · |
| D4 | 0.7310 | #615 | 4 | I | · | P | S | · | · | A | · | · | · | · | · | · | · | · | · | · | · | · | · | · | · |
| D5 | 0.7269 | #1902 | 4 | · | · | P | S | · | L | A | · | · | · | · | · | · | · | · | · | · | · | · | · | · | · |
| Experimental | 0.6074 | — | 20 | I | Q | P | S | S | L | A | T | · | K | K | A | · | S | S | L | L | P | F | T | Q | V |
| Sapiens | 0.6285 | — | 15 | I | Q | P | S | S | L | A | T | · | · | · | A | · | S | S | L | L | · | · | T | Q | · |
| Hu-mAb | 0.4295 | — | 10 | · | Q | · | S | · | · | A | · | R | · | · | A | L | S | · | · | L | P | · | T | · | · |

Aliphatic AVILM · Aromatic FWY · Polar STNQC · Basic KRH · Acidic DE · Gly/Pro GP

**Figure S3. Sequence substitutions in pertuzumab candidates and reference sequences.** Panels A and B show the VH and VL alignments, respectively. Top1–Top5 are the highest-reward unique VH/VL pairs pooled from five independent searches. D1–D5 were selected by greedy farthest-first sampling from pairs with median observed reward ≥98% of the case maximum. Reward is the median across runs in which each pair appeared; ranks refer to the pooled unique pairs. Experimental, Sapiens, and Hu-mAb comparators are unranked. Mutation counts are parent-relative substitutions in each chain. Columns include all mutated Kabat positions across candidates and comparators. Dots indicate parental residues; colors indicate replacement-residue classes; asterisks mark Kabat CDR positions.

**(A)** Bevacizumab · VH

| Candidate / Method | Reward | Rank | Mutations | H1 | H2 | H6 | H9 | H10 | H12 | H13 | H16 | H17 | H18 | H20 | H23 | H38 | H43 | H46 | H48 | H49 | H62* | H63* | H68 | H69 | H72 | H75 | H76 | H82 | H82A | H82B | H83 | H84 | H85 | H89 | H91 | H94 | H105 | H108 |
|---|---|---|---|---|---|---|---|---|---|---|---|---|---|---|---|---|---|---|---|---|---|---|---|---|---|---|---|---|---|---|---|---|---|---|---|---|---|---|
| **Parental** | | | 0 | E | I | Q | P | E | K | Q | E | T | V | I | K | K | K | K | M | G | D | F | T | F | E | A | S | I | S | N | K | N | D | T | F | K | A | T |
| Top1 | 0.7056 | #1 | 11 | · | · | · | · | · | · | K | · | S | · | · | · | R | Q | E | · | · | · | · | · | · | D | · | N | · | N | · | · | · | · | · | Y | R | R | · |
| Top2 | 0.7037 | #2 | 12 | · | · | · | · | · | · | K | · | S | · | · | · | R | Q | E | · | · | · | · | · | · | D | · | N | · | N | · | Q | · | · | · | Y | R | R | · |
| Top3 | 0.7036 | #3 | 12 | · | V | · | · | · | · | K | · | S | · | · | · | R | Q | E | · | · | · | · | · | · | D | · | N | · | N | · | · | · | · | · | Y | R | R | · |
| Top4 | 0.7033 | #4 | 11 | · | · | · | · | · | · | K | · | S | · | · | · | R | Q | E | · | · | · | · | · | · | D | · | N | · | N | · | · | · | · | · | Y | R | R | · |
| Top5 | 0.7032 | #5 | 12 | · | · | · | · | · | · | K | · | S | · | · | · | R | Q | E | · | · | · | · | · | · | D | · | N | · | N | · | Q | · | · | · | Y | R | R | · |
| D1 | 0.7056 | #1 | 11 | · | · | · | · | · | · | K | · | S | · | · | · | R | Q | E | · | · | · | · | · | · | D | · | N | · | N | · | · | · | · | · | Y | R | R | · |
| D2 | 0.6966 | #115 | 13 | · | V | · | · | · | · | K | · | S | · | · | · | R | · | E | · | · | · | · | · | · | D | · | N | · | N | · | R | · | E | · | Y | R | R | · |
| D3 | 0.6950 | #179 | 11 | · | · | · | · | · | · | K | · | S | · | · | · | R | Q | E | · | · | · | · | · | · | D | · | N | · | N | · | · | · | · | · | Y | R | R | · |
| D4 | 0.6928 | #266 | 13 | · | V | · | · | · | · | K | · | S | · | · | · | R | Q | E | · | · | · | · | · | · | D | · | N | · | N | · | Q | · | · | · | Y | R | R | · |
| D5 | 0.6918 | #307 | 11 | · | V | · | · | · | · | · | · | S | · | · | · | R | Q | E | · | · | · | · | · | · | D | · | N | · | N | · | · | · | · | · | Y | R | R | · |
| Experimental | 0.4978 | — | 25 | · | V | E | G | G | V | · | G | S | L | L | A | R | · | E | V | · | · | · | · | · | D | K | · | M | N | S | R | A | E | V | Y | · | Q | L |
| Sapiens | 0.5347 | — | 14 | Q | V | · | S | · | · | · | A | S | · | · | · | R | Q | E | · | · | · | · | V | · | D | · | · | · | · | · | · | A | · | V | · | · | R | L |
| Hu-mAb | 0.4305 | — | 14 | · | · | · | G | G | · | · | G | · | L | · | V | A | · | · | · | S | S | V | · | I | · | K | · | · | · | · | R | · | · | V | · | · | K | · |

**(B)** Bevacizumab · VL

| Candidate / Method | Reward | Rank | Mutations | L7 | L8 | L15 | L20 | L22 | L24* | L41 | L42 | L43 | L44 | L46 | L71 | L72 | L77 | L79 | L83 | L100 | L104 |
|---|---|---|---|---|---|---|---|---|---|---|---|---|---|---|---|---|---|---|---|---|---|
| **Parental** | | | 0 | T | T | L | I | S | S | D | G | T | V | V | Y | S | N | E | I | G | L |
| Top1 | 0.7056 | #1 | 10 | · | P | · | T | T | · | G | · | A | P | L | · | · | · | Q | F | Q | · |
| Top2 | 0.7037 | #2 | 9 | S | · | · | T | T | · | G | · | A | P | L | · | · | · | Q | · | Q | · |
| Top3 | 0.7036 | #3 | 9 | · | P | · | T | T | · | G | · | A | P | L | · | · | · | Q | · | Q | · |
| Top4 | 0.7033 | #4 | 10 | · | · | · | T | T | · | G | K | A | P | L | · | · | · | Q | F | Q | · |
| Top5 | 0.7032 | #5 | 9 | · | P | · | T | T | · | G | · | A | P | L | · | · | · | Q | · | Q | · |
| D1 | 0.7056 | #1 | 10 | · | P | · | T | T | · | G | · | A | P | L | · | · | · | Q | F | Q | · |
| D2 | 0.6966 | #115 | 7 | · | · | V | · | · | · | G | · | A | P | L | · | · | · | Q | · | Q | · |
| D3 | 0.6950 | #179 | 9 | S | · | · | T | T | · | G | K | · | P | · | · | · | · | Q | · | Q | V |
| D4 | 0.6928 | #266 | 7 | · | · | V | · | · | · | G | K | A | P | · | · | · | · | · | F | Q | · |
| D5 | 0.6918 | #307 | 8 | · | · | V | T | · | · | G | · | A | P | · | · | T | · | · | F | Q | · |
| Experimental | 0.4978 | — | 16 | S | P | V | T | T | · | G | K | A | P | · | F | T | S | Q | F | Q | V |
| Sapiens | 0.5347 | — | 15 | S | P | V | T | T | · | G | K | A | P | L | F | T | · | Q | · | Q | V |
| Hu-mAb | 0.4305 | — | 9 | · | P | V | · | · | Q | G | K | A | P | · | F | · | · | Q | · | · | · |

Aliphatic AVILM    Aromatic FWY    Polar STNQC    Basic KRH    Acidic DE    Gly/Pro GP

**Figure S4. Sequence substitutions in bevacizumab candidates and reference sequences.** Panels A and B show the VH and VL alignments, respectively. Top1–Top5 are the highest-reward unique VH/VL pairs pooled from five independent searches. D1–D5 were selected by greedy farthest-first sampling from pairs with median observed reward ≥98% of the case maximum. Reward is the median across runs in which each pair appeared; ranks refer to the pooled unique pairs. Experimental, Sapiens, and Hu-mAb comparators are unranked. Mutation counts are parent-relative substitutions in each chain. Columns include all mutated Kabat positions across candidates and comparators. Dots indicate parental residues; colors indicate replacement-residue classes; asterisks mark Kabat CDR positions.

### (A) Omalizumab · VH

Positions (columns of the alignment): H1 H5 H9 H13 H15 H16 H17 H19 H21 H23 H25 H27 H30 H40 H43 H44 H48 H49 H57* H60* H61* H63* H65* H67 H68 H69 H70 H71 H73 H75 H77 H79 H81 H82 H82C H83 H85 H89 H105 H108

| Candidate / Method | Reward | Rank | Mutations | H1 H5 H9 H13 H15 H16 H17 H19 H21 H23 H25 H27 H30 H40 H43 H44 H48 H49 H57* H60* H61* H63* H65* H67 H68 H69 H70 H71 H73 H75 H77 H79 H81 H82 H82C H83 H85 H89 H105 H108 |
|---|---|---|---|---|
| **Parental** | | | 0 | D Q P K S Q S S A S T Y T F N K M G S N P L N I S V T R T Q Q F K L A T E T A T |
| Top1 | 0.6965 | #1 | 12 | · · · · · · · · T · S G S P · G · · · · · · · V T I S · · · · S · · · · · · K · |
| Top2 | 0.6959 | #2 | 12 | · · · · · · · · T · S G S P · G · · · · · · · V T I S · · · · S · · · · · · K · |
| Top3 | 0.6953 | #3 | 12 | · · · · · · · · T · S G S P · G · · · · · · · V T I S · · · · S · · · · · · K · |
| Top4 | 0.6947 | #4 | 12 | · · · · · · · · T · S G S P · G · · · · · · · V T I S · · · · S · · · · · · K · |
| Top5 | 0.6946 | #5 | 12 | · · · · · · · · T · S G S P · G · · · · · · · V T M S · · · · S · · · · · · K · |
| D1 | 0.6965 | #1 | 12 | · · · · · · · · T · S G S P · G · · · · · · · V T I S · · · · S · · · · · · K · |
| D2 | 0.6897 | #49 | 12 | · · · · · E · · T · S G S P · G · · · · · · · V T · S · · · · S · · · · · · K · |
| D3 | 0.6873 | #87 | 11 | · · · · · · · · T · S G S P · G · · · · · · · V T · S · · · · S · · · · · · K · |
| D4 | 0.6834 | #169 | 9 | · · · · · · · · T · S · S P · G · · · · · · · V · · S · · · · S · · · · · · Q · |
| D5 | 0.6826 | #182 | 12 | · · · · · · · · T · S · S P · G · · · · · · · V T I S · · K · S · · · · · · Q · |
| Experimental | 0.5211 | — | 35 | E V G Q G G · R S A S · · A K G V A T A D V G F T I S · D K T Y Q M L R · V Q L |
| Sapiens | 0.4756 | — | 16 | Q · · · · · T · T · S · · P K G I · · · · · · · · · S V · K · S · · V · A V K · |
| Hu-mAb | 0.4450 | — | 21 | · L G · G G · R · · A · · A A · V S · A D V · F · · S · G · S · · M · · · V R A |

### (B) Omalizumab · VL

| Candidate / Method | Reward | Rank | Mutations | L3 L9 L12 L13 L15 L17 L19 L22 L24* L33* L42 L43 L45 L55* L57 L58 L60 L74 L76 L77 L78 L79 L80 L83 L85 L86 L100 L104 |
|---|---|---|---|---|
| **Parental** | | | 0 | Q A A V L Q A S K M Q P I G E I A N H P V E E A T F A L |
| Top1 | 0.6965 | #1 | 7 | · · · · · · · · · · · · K · G · · · S R · · A · V Y · · |
| Top2 | 0.6959 | #2 | 7 | · · · · · E · · · · · · K · · · · · S R · · A · V Y · · |
| Top3 | 0.6953 | #3 | 7 | · · · · · · · · · · · · Q · G · · · S R · · A · V Y · · |
| Top4 | 0.6947 | #4 | 7 | · · · · · · · N · · · · K · · · · · S R · · A · V Y · · |
| Top5 | 0.6946 | #5 | 7 | · · · · · · · · · · · · K · G · · · S R · · A · V Y · · |
| D1 | 0.6965 | #1 | 7 | · · · · · · · · · · · · K · G · · · S R · · A · V Y · · |
| D2 | 0.6897 | #49 | 7 | · · · · · · · · · · · · K · · · · · S R · · A · V Y Q · |
| D3 | 0.6873 | #87 | 8 | V · · · · · · N · · · · K · · · · · S R · · A · V Y · · |
| D4 | 0.6834 | #169 | 8 | · · · · · E · · · · · · K · G · · · S R · · A · V Y · · |
| D5 | 0.6826 | #182 | 6 | · · · · · · · · · · · · K · · · · · S R · · A V · Y · · |
| Experimental | 0.5211 | — | 25 | · S S A V D V T R · K A K E G V S T S S L Q P F · Y Q V |
| Sapiens | 0.4756 | — | 14 | V · · · · E · · · · · · K · G V D T S R · · A V V Y Q · |
| Hu-mAb | 0.4450 | — | 19 | · S S A V D · T R L K A · Q · V S · · S L Q P F · · · V |

Aliphatic AVILM; Aromatic FWY; Polar STNQC; Basic KRH; Acidic DE; Gly/Pro GP

**Figure S5. Sequence substitutions in omalizumab candidates and reference sequences.** Panels A and B show the VH and VL alignments, respectively. Top1–Top5 are the highest-reward unique VH/VL pairs pooled from five independent searches. D1–D5 were selected by greedy farthest-first sampling from pairs with median observed reward ≥98% of the case maximum. Reward is the median across runs in which each pair appeared; ranks refer to the pooled unique pairs. Experimental, Sapiens, and Hu-mAb comparators are unranked. Mutation counts are parent-relative substitutions in each chain. Columns include all mutated Kabat positions across candidates and comparators. Dots indicate parental residues; colors indicate replacement-residue classes; asterisks mark Kabat CDR positions.

### (A) Eculizumab · VH

| Candidate / Method | Reward | Rank | Mutations | H5 | H11 | H12 | H20 | H25 | H28 | H37 | H38 | H40 | H43 | H48 | H50* | H64* | H66 | H67 | H68 | H69 | H71 | H75 | H76 | H78 | H81 | H82B | H83 | H87 | H105 | H108 |
|---|---|---|---|---|---|---|---|---|---|---|---|---|---|---|---|---|---|---|---|---|---|---|---|---|---|---|---|---|---|---|
| **Parental** | | | 0 | Q | L | M | M | T | I | I | K | R | H | I | E | K | K | A | A | F | A | S | N | A | Q | S | T | S | A | T |
| Top1 | 0.6883 | #1 | 12 | · | · | K | · | S | T | V | R | · | · | M | · | · | · | · | T | M | R | · | · | · | E | · | R | · | R | · |
| Top2 | 0.6863 | #2 | 12 | · | · | K | · | S | T | V | R | · | · | M | · | · | · | · | T | M | R | · | · | · | E | · | R | · | R | · |
| Top3 | 0.6863 | #3 | 12 | · | · | K | · | S | T | V | R | · | · | M | · | · | · | · | T | M | R | · | · | · | E | · | R | · | R | · |
| Top4 | 0.6863 | #4 | 11 | · | · | K | · | S | T | V | R | · | · | M | · | · | · | · | · | M | R | T | · | · | · | · | R | · | R | · |
| Top5 | 0.6858 | #5 | 12 | · | · | K | · | S | T | V | R | · | · | M | · | · | R | · | · | M | R | T | · | · | · | · | R | · | R | · |
| D1 | 0.6883 | #1 | 12 | · | · | K | · | S | T | V | R | · | · | M | · | · | · | · | T | M | R | · | · | · | E | · | R | · | R | · |
| D2 | 0.6832 | #41 | 11 | · | · | K | · | S | T | V | R | · | · | M | · | · | · | · | · | M | R | T | · | · | · | · | R | · | R | · |
| D3 | 0.6816 | #96 | 13 | · | · | K | · | S | T | V | R | · | · | M | · | · | R | · | · | M | R | T | · | · | · | G | R | · | R | · |
| D4 | 0.6762 | #608 | 13 | · | V | K | · | S | T | V | R | · | · | M | · | · | · | · | T | M | R | · | · | · | E | · | R | · | R | · |
| D5 | 0.6762 | #612 | 12 | · | · | K | · | S | T | · | R | · | Q | M | · | · | · | · | T | M | R | · | · | · | E | · | R | · | R | · |
| Experimental | 0.5036 | — | 23 | V | V | K | V | S | · | V | R | A | Q | M | · | · | R | V | T | M | R | T | S | V | E | · | R | T | Q | L |
| Sapiens | 0.4663 | — | 15 | V | V | K | I | S | · | V | R | A | Q | · | · | · | R | V | · | · | · | · | · | · | E | · | · | T | R | L |
| Hu-mAb | 0.3501 | — | 15 | V | V | K | V | · | · | · | R | A | · | M | W | Q | R | V | · | · | · | I | · | · | E | · | · | T | C | · |

### (B) Eculizumab · VL

| Candidate / Method | Reward | Rank | Mutations | L9 | L17 | L18 | L24* | L38 | L40 | L43 | L45 | L55* | L58 | L59 | L69 | L70 | L71 | L72 | L74 | L79 | L81 | L83 | L100 | L104 | L106 |
|---|---|---|---|---|---|---|---|---|---|---|---|---|---|---|---|---|---|---|---|---|---|---|---|---|---|
| **Parental** | | | 0 | A | E | T | G | R | Q | S | Q | A | M | S | R | Q | Y | Y | K | H | D | V | A | L | L |
| Top1 | 0.6883 | #1 | 8 | · | · | · | · | · | · | A | K | · | V | P | T | · | · | T | T | · | · | · | · | V | · |
| Top2 | 0.6863 | #2 | 8 | · | · | R | · | · | · | A | K | · | V | · | T | · | · | T | T | · | · | · | Q | · | · |
| Top3 | 0.6863 | #3 | 8 | · | D | · | · | · | · | A | K | · | V | P | T | · | · | T | T | · | · | · | · | · | · |
| Top4 | 0.6863 | #4 | 9 | · | · | · | · | · | P | · | K | · | V | P | · | D | · | T | T | Q | · | · | · | · | I |
| Top5 | 0.6858 | #5 | 8 | · | D | · | · | · | P | · | K | · | V | · | · | D | · | T | T | Q | · | · | · | · | · |
| D1 | 0.6883 | #1 | 8 | · | · | · | · | · | · | A | K | · | V | P | T | · | · | T | T | · | · | · | · | V | · |
| D2 | 0.6832 | #41 | 9 | · | D | · | · | Q | P | · | · | · | V | · | · | D | · | T | T | Q | · | · | Q | · | · |
| D3 | 0.6816 | #96 | 7 | · | · | · | · | · | P | · | K | · | V | · | · | D | · | T | T | Q | · | · | · | · | · |
| D4 | 0.6762 | #608 | 5 | · | · | · | · | · | · | A | K | · | · | · | · | · | F | T | T | · | · | · | · | · | · |
| D5 | 0.6762 | #612 | 5 | · | · | · | · | Q | · | · | · | · | · | · | T | · | · | T | T | · | · | · | · | · | I |
| Experimental | 0.5036 | — | 20 | S | D | R | · | Q | P | A | K | · | V | P | T | D | F | T | T | Q | E | F | Q | V | I |
| Sapiens | 0.4663 | — | 14 | S | D | · | · | Q | P | A | · | · | V | · | T | D | F | T | T | Q | · | · | Q | · | I |
| Hu-mAb | 0.3501 | — | 12 | S | D | R | R | · | P | A | · | Q | V | · | T | · | F | · | T | · | · | · | Q | · | · |

Aliphatic AVILM | Aromatic FWY | Polar STNQC | Basic KRH | Acidic DE | Gly/Pro GP

**Figure S6. Sequence substitutions in eculizumab candidates and reference sequences.** Panels A and B show the VH and VL alignments, respectively. Top1–Top5 are the highest-reward unique VH/VL pairs pooled from five independent searches. D1–D5 were selected by greedy farthest-first sampling from pairs with median observed reward ≥98% of the case maximum. Reward is the median across runs in which each pair appeared; ranks refer to the pooled unique pairs. Experimental, Sapiens, and Hu-mAb comparators are unranked. Mutation counts are parent-relative substitutions in each chain. Columns include all mutated Kabat positions across candidates and comparators. Dots indicate parental residues; colors indicate replacement-residue classes; asterisks mark Kabat CDR positions.

### Figure S7

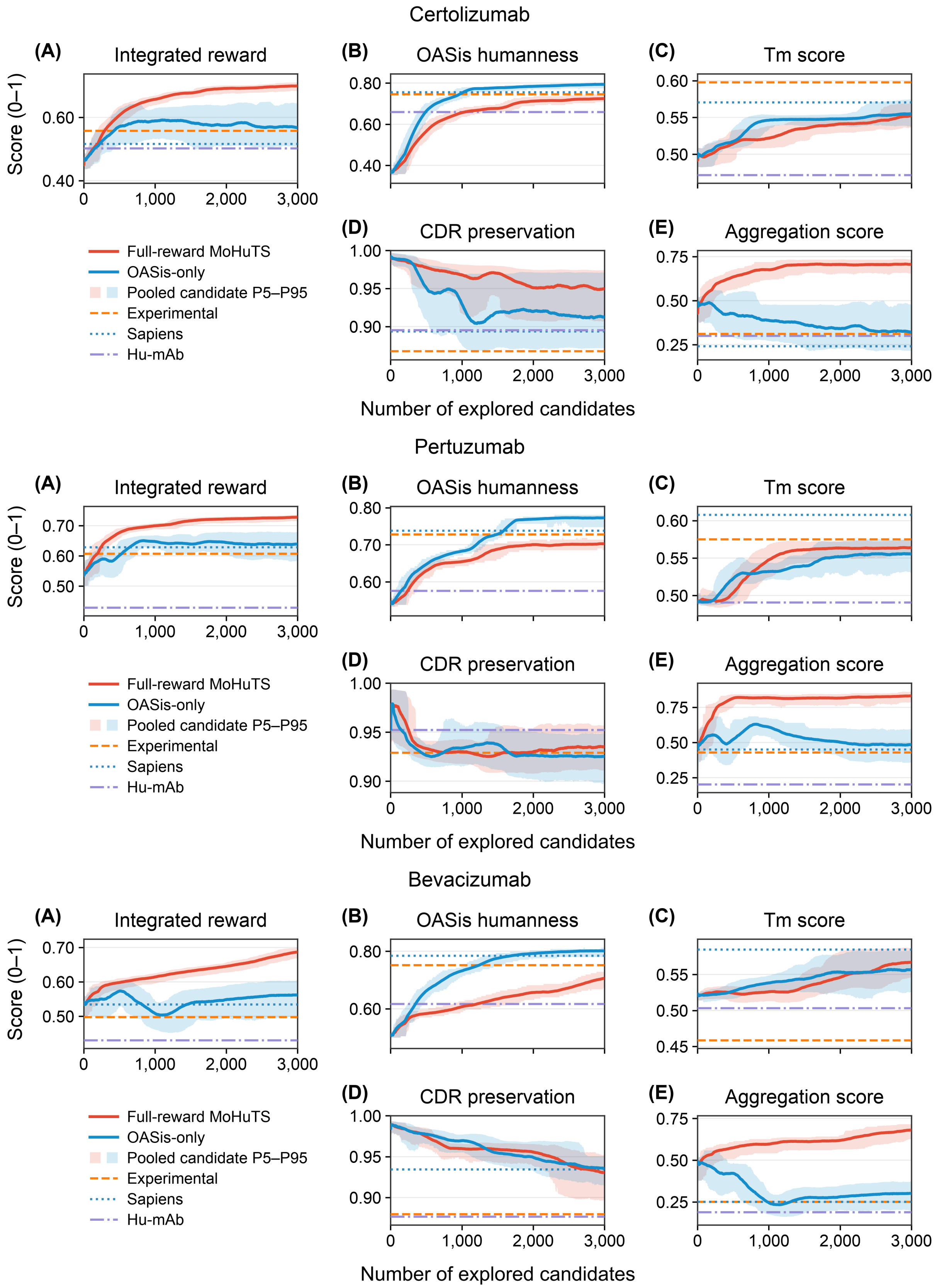

Certolizumab
(A) Integrated reward
(B) OASis humanness
(C) Tm score
(D) CDR preservation
(E) Aggregation score
Score (0–1)
Number of explored candidates
Full-reward MoHuTS
OASis-only
Pooled candidate P5–P95
Experimental
Sapiens
Hu-mAb
Pertuzumab
(A) Integrated reward
(B) OASis humanness
(C) Tm score
(D) CDR preservation
(E) Aggregation score
Score (0–1)
Number of explored candidates
Full-reward MoHuTS
OASis-only
Pooled candidate P5–P95
Experimental
Sapiens
Hu-mAb
Bevacizumab
(A) Integrated reward
(B) OASis humanness
(C) Tm score
(D) CDR preservation
(E) Aggregation score
Score (0–1)
Number of explored candidates
Full-reward MoHuTS
OASis-only
Pooled candidate P5–P95
Experimental
Sapiens
Hu-mAb

**Figure S7 (continued)**

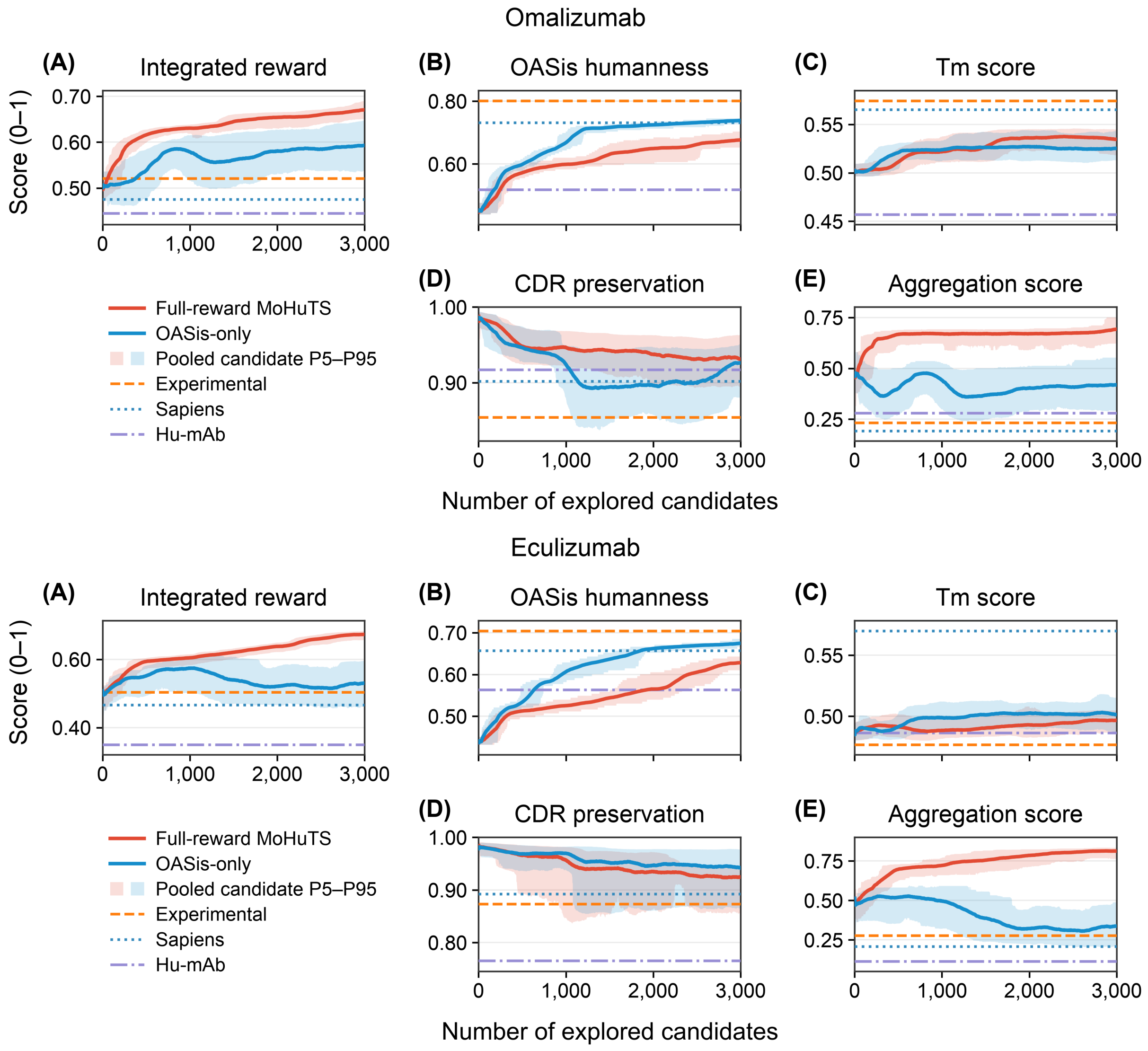


**Figure S7.** Full-reward and OASis-only search trajectories for five antibody cases: certolizumab, pertuzumab, bevacizumab, omalizumab, and eculizumab (top to bottom). Panels A–E show integrated reward, OASis humanness, normalized predicted Tm, CDR preservation, and aggregation scores, respectively. Red and blue lines show full-reward MoHuTS and OASis-only searches, respectively ($n = 5$ per condition and antibody). Each run explored 3,000 unique VH/VL sequences, with matched seeds and settings except reward weights. All candidates were evaluated using the same four normalized components; integrated rewards for OASis-only candidates were calculated retrospectively. Lines show trailing 200-candidate means averaged across five runs; means for the first 199 candidates use all candidates explored so far per run. Shading shows the 5th–95th percentiles pooled across runs within these windows, reflecting candidate variation, not confidence intervals across runs. Horizontal lines show experimentally validated humanized (Experimental), Sapiens, and Hu-mAb references. Higher scores are more favorable. The x-axis shows unique candidates explored per run.
